# Magnetic DNA random access memory with nanopore readouts and exponentially-scaled combinatorial addressing

**DOI:** 10.1101/2021.09.15.460571

**Authors:** Billy Lau, Shubham Chandak, Sharmili Roy, Kedar Tatwawadi, Mary Wootters, Tsachy Weissman, Hanlee P. Ji

## Abstract

The storage of data in DNA typically involves encoding and synthesizing data into short oligonucleotides, followed by reading with a sequencing instrument. Major challenges include the molecular consumption of synthesized DNA, issues with basecalling errors, and limitations with scaling up read access operations for individual data elements. Addressing these challenges, we describe a DNA storage system called MDRAM (**M**agnetic **D**NA-based **R**andom **A**ccess **M**emory) that enables repetitive and efficient readouts of targeted files with nanopore-based sequencing. Through conjugation of synthesized DNA to magnetic beads, we enabled repeated readouts of data while preserving the original DNA analyte and maintaining data readout quality. MDRAM also utilizes an efficient convolutional coding scheme that leverages soft information in raw nanopore sequencing signals to achieve information reading costs comparable to Illumina sequencing despite substantially higher error rates. Finally, we demonstrate a proof-of-concept DNA-based proto-filesystem that enables an exponentially-scalable data address space using only small numbers of targeting primers for assembly and readout.

**ONE-SENTENCE SUMMARY:** We demonstrate a novel DNA data storage system that leverages conjugation of DNA onto magnetic beads, new computational advances in data encoding, and exponentially scalable access of individual data elements.

## INTRODUCTION

DNA has properties that allow it to store and maintain genetic information for extended periods of time. As a result, DNA provides a potential solution to the need for massive data storage. DNA molecules provide extremely high data density and long-term durability (*1, 2*). In comparison to current magnetic and solid-state data storage technologies, DNA has a potentially higher storage density in addition to natural redundancy from molecular copying. Magnetic and solid-state storage technologies can be stably maintained up to decades but undergo progressive media degradation. In contrast, under optimal storage conditions, the physical stability of DNA is on the order of thousands of years (*1, 2*). There is ongoing work to develop DNA storage technologies with data writing into encoded oligonucleotides and reading through next generation sequencers (*3*).

Data storage systems revolve around the concept of random access of data elements from a larger stored pool of multiple data stores or ‘files’. New molecular technologies will be required for recovering stored data in synthetic DNA on an arbitrarily large scale with random access features. Earlier studies used wholesale sequencing of the entire archive followed by bioinformatic extraction of small parts of data from the dataset (*4, 5*). Recently, some groups have used conventional PCR to amplify the target files of interest for sequence-based reading (*6-8*). However, this method has limitations on its scalability to the large numbers of DNA-based data files. For DNA-based data, reading files typically requires PCR amplification with individual pairs of targeting primer oligonucleotides. Accessing these files also leads to consumption and potential representative distortions of the DNA molecular information. These issues are a direct result of the molecular preparation and amplification for the sequencing process. Recent work has shown it is possible to re-amplify DNA oligonucleotide pools (*9*) but an ideal solution would be potentially lossless reading of a file from the source DNA.

Errors in DNA synthesis, artifacts in sequencing, and sequence dropouts also occur frequently. As a result, reading DNA data requires error correction techniques to enable reliable recovery of data and to optimize the coding density and sequencing coverage. Erlich and Zielinski focused on Illumina sequencing with a Fountain code scheme (*9, 10*). Organick et al. presented a scheme for both Illumina and nanopore sequencing using a variety of methods including multiple sequence alignment (MSA) and Reed Solomon codes (*11*) to handle sequence erasure and sequencing errors (*7*). Lopez et al. improved the MSA algorithm leading to significantly fewer reads being required for successful decoding with nanopore sequencing (*12*). Several theoretical studies focused on various aspects of the DNA-based storage problem such as the information-theoretic capacity in the asymptotic setting (*13*), the optimality of various techniques to recover the order of the oligonucleotides (*14*), development of indel correction codes (*15*) and the tradeoff between the writing and reading cost associated with DNA storage (*16*). Nanopore sequencing is also subject to unique error profiles consisting of substantially higher rates of indels. Because of the higher error rates in nanopore sequencing, developing a robust data encoding scheme is a requirement when using this platform for DNA data reading.

Addressing the challenges of error correction and iterative random data access, we developed a highly scalable random access file system called magnetic DNA-based random access memory **(MDRAM)**. This technology uses new advances in DNA encoding for maximum reading efficiency and repeated data readout operations. MDRAM has several fundamental innovations in the field of DNA data storage **(Fig. 1A)**. This system involves conjugating arbitrary DNA data files, in the form of oligonucleotides, to solid-phase magnetic beads. DNA data files can be selectively sequenced without losing the original molecular DNA media – this feature enables multiple repetitive read operations. We improve DNA encoding using convolutional codes that decrease the reading costs of current methods by an order of magnitude. This feature leverages rapid nanopore sequencing readouts and thus enables low-cost, real-time data retrieval. Using MDRAM we also demonstrate a combinatorial barcoding system for labeling and retrieving data packets that can scale exponentially to potentially unlimited numbers of files. This enables a unique DNA targeting scheme that is exponentially scalable without the burden of individually synthesizing individual primer pairs for enrichment. Overall, we demonstrate significant improvements towards the implementation of robust encoding and scalable access of DNA elements.

**Figure 1.**
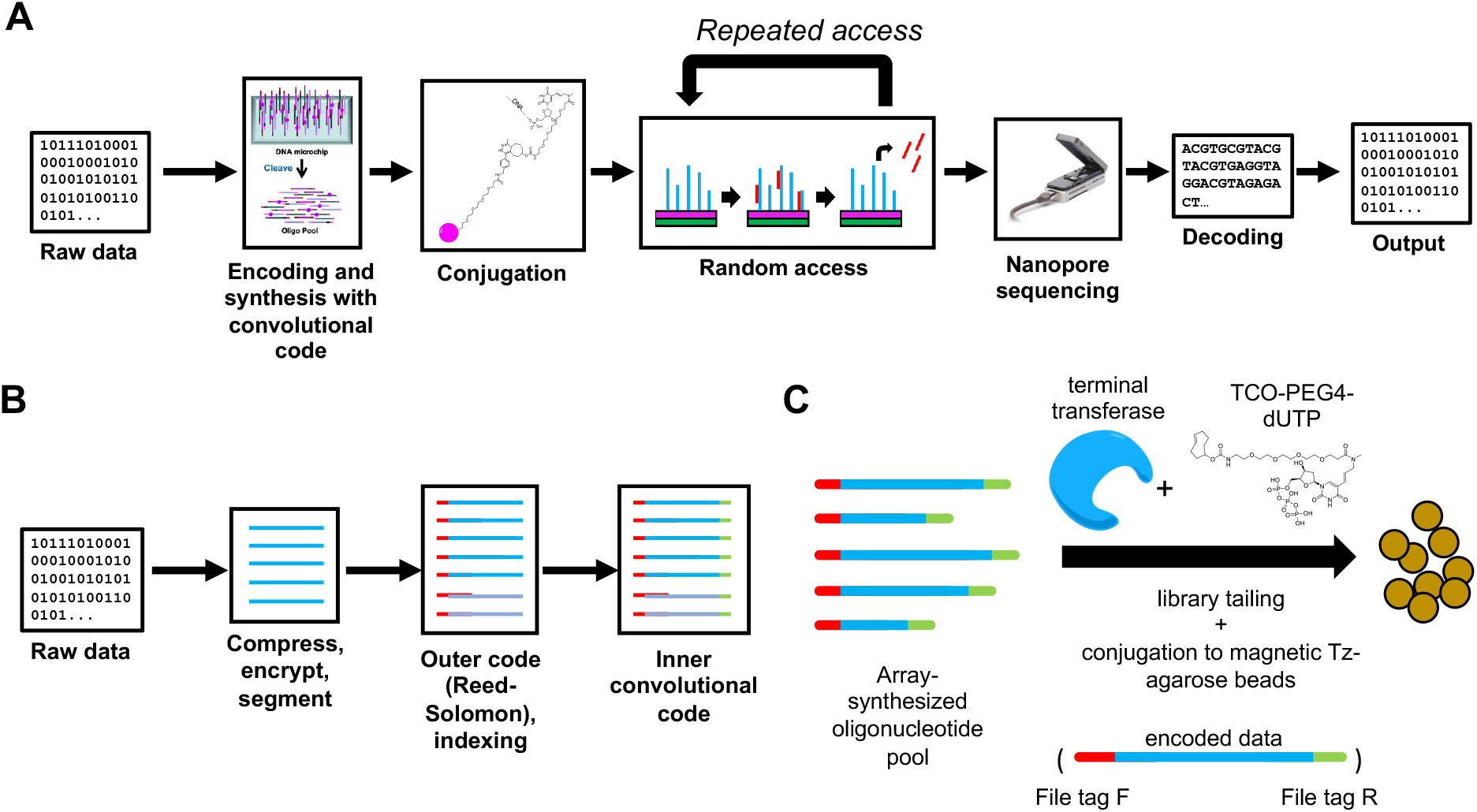
MDRAM Overview. (A) Raw data is encoded into nucleotide bases using a convolutional encoding workflow, and are synthesized as an oligonucleotide pool. These fragments are then conjugated onto functionalized magnetic agarose beads, after which specific data elements can be repeatedly targeted by PCR. These fragments can be sequenced on the Oxford Nanopore system, after which they are decoded using an integrated workflow that does not rely on individual basecalled nucleotides. (B) Data encoding. Information is encoded with both an outer and inner code. The outer code (Reed-Solomon) accounts for oligonucleotide dropouts that may occur during synthesis. The inner code accounts for errors that may occur during the synthesis and sequencing processes. (C) Array-synthesized oligonucleotides are functionalized with *trans-*cyclooctene dUTP with terminal transferase, and then conjugated to methyltetrazine-functionalized magnetic agarose beads. Each oligonucleotide contains adapter sequences (red and green) such that subpools of oligonucleotides can be enriched by PCR.

## RESULTS

### Encoding DNA with convolutional codes

In this study, we designed and optimized a schema for writing data into a DNA information payload with a convolutional encoding method optimized for nanopore sequencing (*17*). Designed for use with nanopore sequencing, encoded oligonucleotides contain an inner code to mitigate sequencing errors and a Reed Solomon (RS) outer code (*11*) to mitigate sequence dropouts **(Fig. 1B)**. We employed convolutional coding as the inner code. A convolutional encoder encodes a stream of message bits into a sequence of encoded bits, which are computed as a linear combination of a past window of m input bits (*17*). The rate parameter r refers to the coding rate, which is the ratio of input to output encoded bits. As an example of how these two parameters influence DNA data encoding and decoding, let us consider an example where the convolutional code has the following value of m=6 and r=0.5; this produces 2 output bits per input bit. As m increases, the code becomes more powerful, but the decoding becomes slower due to an exponential increase in the number of possible decoded states. Moreover, each data sequence has a convolutional code and a cyclic redundancy check **(CRC)** error-detecting code (*18*). Additional sequence elements incorporated in the synthesized oligonucleotides also consist of adapter sequences for targeted PCR amplification, which flank the internal index sequence and encoded data payload **(Supplementary Figure S1)**.

To test the chemistry underlying MDRAM media, we used an oligonucleotide pool denoted as Pool A. This set of contains ∼12,000 unique array-synthesized oligonucleotides spanning 13 subpools (‘files’) that can be targeted by PCR. The sequences encoded a collection of various song lyrics, speeches, and the Universal Declaration of Human Rights totaling a size of 11.3 KB. Each of the 13 subpools encoded the same compressed archive of the aforementioned material using an early implementation of the convolutional encoding scheme, using a variety of m and r parameters; Pool A’s encoding performance was described previously (*17*).

### Stable retention of DNA data elements onto magnetic beads

Encoded DNA was functionalized by adding *trans*-cyclooctene **(TCO)** modified dUTPs with terminal transferase. This enzymatic step enables the conjugation onto customized methyltransferase **(Tz)** functionalized magnetic agarose beads – the chemistry involves an inverse electron demand Diels-Alder cycloaddition **(Fig. 1C)**. Amongst all variations of click chemistry schemes, the kinetic rate of TCO-Tz conjugation is particularly suitable for modifying molecular analytes at submicromolar concentrations (*19*).

First, we evaluated the efficiency of the conjugation chemistry with a variety of experiments. This step involved conjugation reactions with 70 nanograms of Pool A by adding TCO-functionalized dUTPs via terminal transferase (*20*) and then mixing the product with Tz-functionalized magnetic beads. After the conjugation reaction, we separated and collected the supernatant of the original reaction. We used fluorimetry to quantify the amount of remaining DNA left in the supernatant after the conjugation reaction. The amount of remaining DNA was below the limit of detection, indicating that the majority of DNA was conjugated onto the magnetic beads. To verify this fluorimetry result, we performed qPCR on the supernatant of the conjugation reaction and the eluent from different washings of the MDRAM beads. As a measure of residual DNA material left in the supernatant and washes, we quantified the Ct value corresponding to a single file from the oligonucleotide pool and used a serial dilution of the oligonucleotide pool stock as the quantitative standard. Less than 0.01% DNA material was detected in the conjugation supernatant and in each wash step **(Fig. 2A)**. These results indicated a near complete, stable conjugation of the DNA material to the magnetic agarose beads. This high level of efficiency is consistent with other published measurements of Tz-TCO’s conjugation efficiency with biomolecules (*19*).

**Figure 2.**
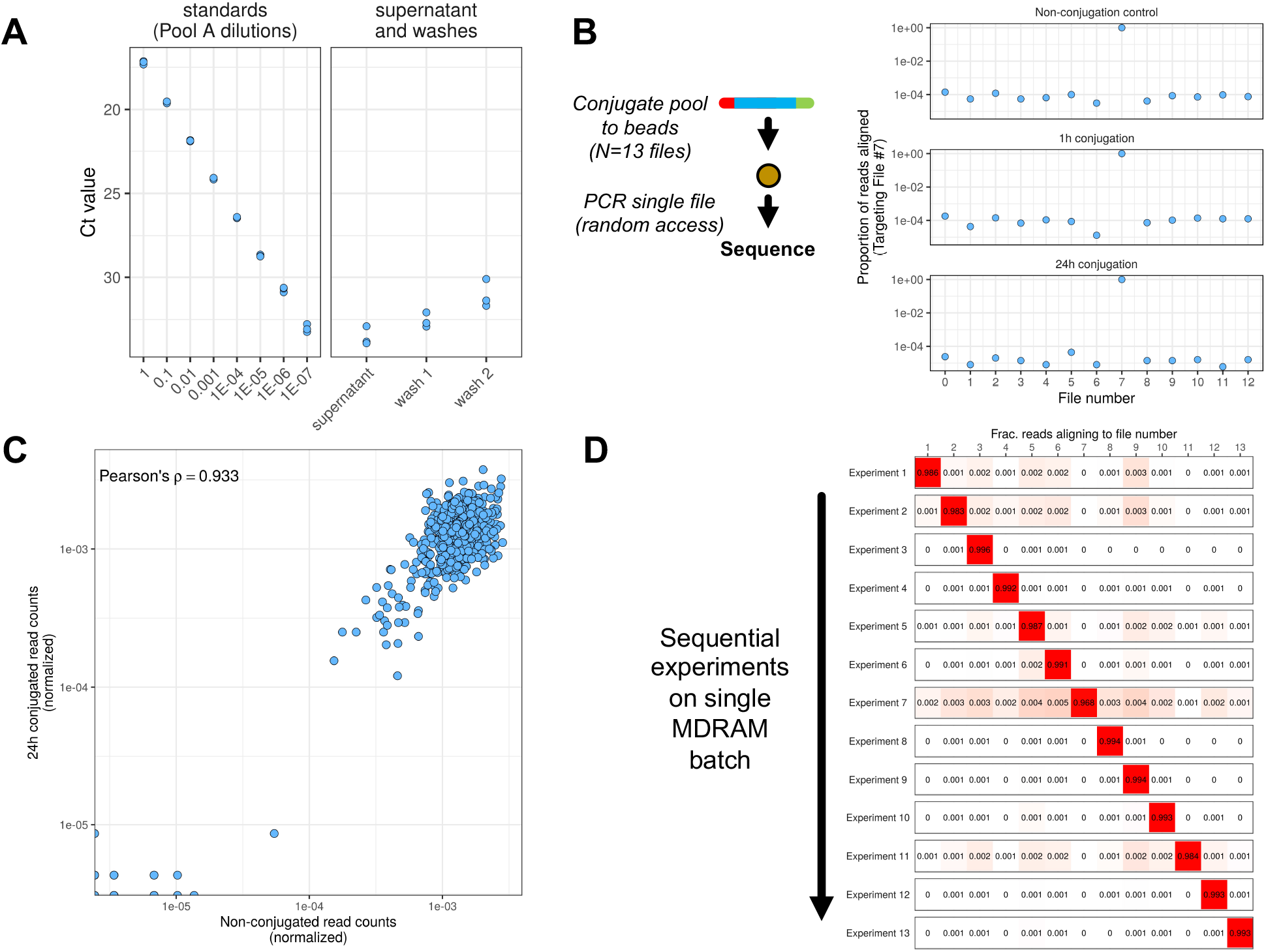
Random access of data elements with MDRAM. (A) Assessment of conjugation efficiency. qPCR was performed on the supernatant, which measures the amount of DNA not conjugated to the magnetic beads and after wash steps. Left: Standard curve of DNA ranging several orders of magnitude. Right: quantitation of residual DNA from the conjugation reaction and through wash steps. Each dot represents a qPCR replicate. (B) Random access of a data element. A single DNA data file was amplified from MDRAM with different conjugation times (1 hour and 24 hours), and its on-target alignment rate was compared to an unconjugated control experiment with an aqueous DNA template. (C) Assessment of oligonucleotide abundances. The distribution of individual oligonucleotides from a targeted file was measured against the unconjugated control. The Pearson correlation coefficient was 0.933. (D) Iterative and sequential random access of data elements. Individual data files were sequentially retrieved from MDRAM. In between each experiment, the beads were washed, after which another file was targeted. The on-target alignment rate is shown for each random access experiment on the same bead batch.

### Retrieval of data elements from MDRAM

To demonstrate random data access capability, we performed targeted sequencing of specific data files from the MDRAM substrate containing Pool A. To assess the overall performance of DNA amplification from MDRAM we used sequencing to determine the on-target rates of a single DNA data file (e.g. number 7) **(Fig. 2B)**. Two different bead conjugation conditions were tested; one involving a one hour incubation and the other for 24 hours. We performed PCR reactions across two different bead conjugation conditions, sequenced the resultant amplicons on the Illumina platform and counted reads using the entire multi-file oligonucleotide pool as the alignment reference **(Fig. 2B, Supplementary Table S1)**. Illumina-based sequences were used to accurately assess the count distribution of the amplified individual oligonucleotides. As a control, we compared the MDRAM read counts versus those amplified from a baseline in-solution oligonucleotide pool from the original array synthesis. Referred to as the non-conjugation control, this comparison sample involved direct sequencing from PCR amplification of the aqueous oligonucleotide pool. This control did not use the MDRAM media.

We aligned all sequenced reads using the designed oligonucleotide sequences as a reference. The proportion of on-target reads were over 95%. We noted a similar performance between two different conjugation times when compared to non-conjugated controls **(Fig. 2B)**. This result confirmed that reading specific data files is done with high efficiency when the template DNA is conjugated to magnetic agarose beads. We measured the count distribution of detected oligonucleotides against a non-conjugated control and determined the concordance with MDRAM-amplified product. We observed a Pearson correlation coefficient of 0.933. This result indicates that read distributions were not significantly affected when conjugated to magnetic agarose beads versus the in-solution oligonucleotides **(Fig. 2C)**.

### Sequential random access of DNA data elements with nanopore sequencing

MDRAM is a stable storage medium that enables sequential reading of different data files from the same substrate. Using PCR amplification, we performed serial random file access operations for each of the 13 files of Pool A on the same MDRAM media. For this test, we conjugated 70 nanograms of a DNA data oligonucleotide pool A to a new batch of magnetic beads. An Oxford MinION instrument was used to sequence the amplicons. Between each experiment, the beads were washed before PCR amplification for the next file was performed **(Methods)**.

The sequences were aligned to a reference representing the synthesized oligonucleotide sequences. The on-target rate of accessing each file was over 98% for all sequential read operations with each PCR primer pair exhibiting a minimum observed on-target rate of 96.8% **(Fig. 2D, Supplementary Table S2)**. We measured the off-target sequences (i.e. reads aligning to file 1 when targeting for file 2). There was less than 0.1% of crossover contamination from the other DNA file products across all experiments. This result indicates that MDRAM can be used for sequential read operations of specific data files without contaminating reads from other files or previous read iterations.

### Accurate convolutional decoding integrated into nanopore basecalling

To improve the accuracy of reading MDRAM files with nanopore sequence, we developed a decoding workflow that can be integrated with open-source nanopore basecaller software **(Methods)** (*17*). For this study, we used the Bonito basecaller (*21*), which processes nanopore signal data using neural networks. In this approach, information is drawn directly from the nanopore raw electrical current signals rather than from basecalled reads – the latter contains artifactual substitutions, insertions, and deletions in the sequences, some of which arise because of nanopore signal fluctuations. Convolutional decoding (*22*) enables serial error correction defined by a state transition diagram; the decoding is thus performed efficiently using a dynamic programming-based Viterbi algorithm. In the Bonito basecaller framework, upstream of the basecalls are probability scores that can be integrated with state transitions in the convolutional decoding framework. The output is then a list of top candidate codewords, denoted as L and a default value of 8. Additional filtering comes from the CRC technique (*18*) which provides added error detection and allows one to discard reads under circumstances where the convolutional code makes an error. Full details and source code are described in **Methods**.

We measured the performance of our convolutional encoding and decoding scheme across multiple encoding parameters. We used an oligonucleotide pool (‘Pool B’) with approximately 15,000 non-overlapping sequences. Pool B had an expanded number of data files compared to Pool A, with a 12.7 KB dataset used for each parameter setting. Pool B focused on higher rate and lower memory codes to test the limits of convolutional coding, and to move towards practical systems with lower writing costs and computational requirements **(Supplementary Table S3)**. Also, Pool B incorporated additional encoding strategies and was synthesized using high-fidelity array synthesis chemistry from Agilent (*23, 24*). Details of the additional files are listed in **Methods**. Each data file, corresponding to an oligonucleotide subpool, was PCR amplified with primers flanking the file of interest **(Supplementary Table S4)** and underwent nanopore sequencing **(Supplementary Table S2)**.

We analyzed the decoding performance as influenced by two encoding parameters, the convolutional code memory (m) and rate (r) on a random subsampling of reads (N=10 trials). We assessed the percent of successfully decoded reads for the different values of m and r, meaning whether the sequenced data successfully decoded back to the file of interest **(Supplementary Table S5)**. As expected, we observed that a higher memory and lower rate consistently led to a greater fraction of successfully decoded reads (up to 88%) at the cost of more computation and higher writing cost **(Supplementary Figure S4)**. Interestingly, the results were not monotonic; there was a general trend with codes having a m=11 and with r=3/4 associated with a lower reading cost. The overall improvements were substantial, leading to more than two times higher decoding success rate in some cases **(Supplementary Table S6)** when compared to our prior results (*17*). The improvement is more pronounced for higher rate and lower memory codes; these settings enable the use of more practical and efficient codes while achieving similar performance to more computationally intensive and lower density codes. Across the different subpools, we also explored other optimization strategies, such as utilizing multiple CRC segments and finetuning the Bonito basecaller model but did not observe significant nor consistent improvements in decoding performance **(Supplementary Figure S5, Supplementary Text)**.

We measured other performance metrics that included: writing cost in terms of bases synthesized per information bit; reading cost measured in bases sequenced per information bit; and minimum required read coverage associated with each experimental parameter. Prior reports of DNA data storage (*7, 9*) described the coding density (inverse of writing cost) and coverage (reading cost divided by writing cost). We converted these metrics to the ones mentioned above for comparison **(Methods)**. Previously, we described (*16*) that reading cost is a better representation of the actual cost of sequencing compared to coverage, especially when comparing schemes with unequal writing cost or coding density. We considered the well-performing m=11 codes across all rates (r) alongside selected rates for m=6 and m=8 to simplify the analysis **(Fig. 3A)**. Consistent with the results measuring the proportion of correctly decoded reads, we observed significantly improved reading costs with our convolutional encoding/decoding scheme. Specifically, there were 2-3x lower reading costs compared to our previous work with nanopore sequencing (*17*). We achieved a reading cost of 3-5 bases/bit, compared to 22 and 34 bases/bit achieved by previously published strategies (*7, 12*) for nanopore sequencing based readout. These results indicated a cost that was 10 times lower than other encoding frameworks – we attribute this to reduced reliance on consensus for error correction that was used by the other methods **(Fig. 3A)**.

**Figure 3.**
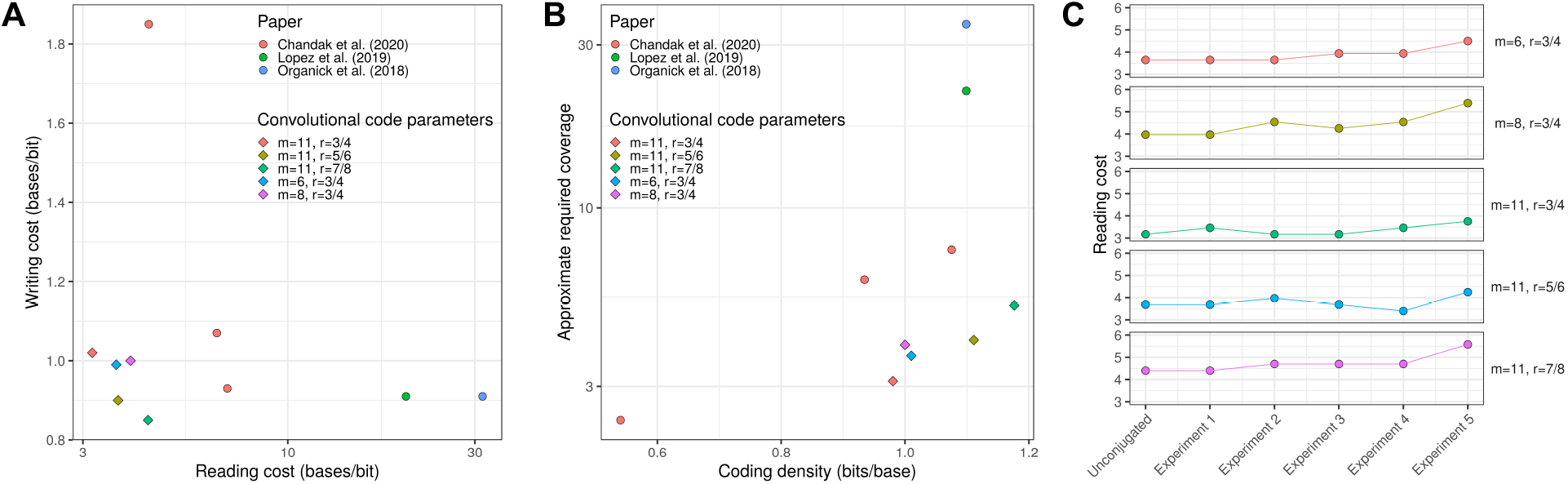
Efficient encoding of information in DNA with convolutional codes. (A) Convolutional code performance. The reading and writing cost of the convolutional code was measured for a variety of parameters and compared to other works. (B) Approximate required sequencing coverage. The amount of sequencing reads in order to successfully decode a file is shown as a function of the coding density (bits/base) and compared to other works. (C) Application of convolutional codes to parallel file access in MDRAM. Multiple files were accessed by MDRAM in parallel by splitting beads and performing PCR. Beads were then pooled together, washed, and split again for another experiment. The reading cost is plotted as a function of experiment number.

For successful decoding in other frameworks, approximately 30-fold coverage is needed (*7*). In contrast, our strategy required as low as 3-fold read coverage from nanopore-based sequencing **(Fig. 3B)**. Most striking is that for writing costs of one base/bit, our nanopore-based framework not only improved on our previous work (*16*), but also is competitive with other works (*7, 9*) that developed coding methods for Illumina sequencing **(Supplementary Table S8)**. This improvement with nanopore sequencing was notable given its substantially lower basecalling accuracy (∼5-10%) compared to Illumina sequencing (∼0.1-1%). This result also points to the strength of convolutional coding and the basecaller-decoder integration framework.

### Repeatable, parallel, and multi-file access on magnetic beads

We leveraged our improved encoding scheme for highly efficient and parallel random access operations on magnetic beads. MDRAM enables multiple parallel random read operations by splitting the bead media. Afterwards, the bead substrate is washed and combined for subsequent data access operations. To demonstrate MDRAM’s performance over multiple iterations, we conducted random access reading on specific subpools. Here, we conjugated 100 nanomoles of Pool B onto magnetic beads. We split the magnetic beads into five wells and in each well we targeted a different file using a paired PCR primer pair specific to that data file **(Supplementary Table S2)**. After amplification, the beads were pooled back together and washed for a new experiment.

We assessed the performance of integrating our improved convolutional coding onto MDRAM beads. This part of the study involved included a non-conjugated control – the original aqueous oligonucleotide pool – for every targeted file of interest in every flowcell to control for run-to-run variation in sequencing quality. Base quality variation was apparent among the different sequencing runs with the mean basecalling error rate varying from 4.5% to 7%. However, there were no significant differences in base quality between the MDRAM versus non-conjugated libraries when sequenced in the same flowcell **(Supplementary Table S7A)**.

We also evaluated whether there were substantial decoding impacts on the MDRAM platform. The outer Reed Solomon code requires a specific minimum number of unique sequences to be correctly decoded. In cases with significant coverage variance or dropout of certain sequences, a higher number of reads is required for decoding the data. We determined that the read coverage variance was higher for the MDRAM-based experiments, leading to a higher reading cost required compared to the aqueous oligonucleotide pool template control **(Supplementary Table S7B, C)**. However, the convolutional code parameters with more powerful error correction capabilities robustly controlled this effect. The code with highest memory (m=11) and lowest rate (r=3/4) had only a 15% variation in reading cost across repeated data readout operations **(Supplementary Table S7D**,**E)**. The decoding performance across iterative read operations had less than 10% variation in the proportion of successfully decoded reads **(Supplementary Table S7F)**. When using MDRAM, the reading cost across iterative random access operations remained up to an order of magnitude lower compared to recent nanopore-based DNA storage methods **(Fig. 3C, Supplementary Table S8)**. Overall, when using MDRAM for repeated access operations, the reading cost of our new encoding framework remained below 5 bases/bit in all but two data points, and below 4 bases/bit for the m=11 and r=3/4 encoding parameters.

### Exponential-scale hierarchical file access

We developed a strategy for improved data access that is scalable to large numbers of data file elements with MDRAM. Read operations that access data elements using conventional PCR are severely bottlenecked by the design and synthesis of individual primer pairs to amplify and read a given file of interest. In a conventional random access scheme where individual data elements are accessed by a PCR amplification, the number of primers scales directly with the number of files. In conventional computer-based file systems, there may be millions of files that are all accessed individually. In a DNA-based storage system, the conventional PCR-based approach for selectively amplifying specific files is restricted in its scalability. It is nearly impossible to synthesize the number of PCR primers required to sequencing all the various files. Another challenge is that a significant number of primer pairs will have off-target amplification for any given file of interest.

Providing a solution for this constraint, we improved the read operation process with a hierarchical file system for MDRAM. We refer to this scheme as combinatorial barcode addresses **(CBAs)**. With it, an exponentially scalable set of different data elements can be tagged and accessed with only a small number of oligonucleotide sequences **(Fig. 4A)**. CBAs use a highly diverse combinatorial schema for sequence tagging and retrieval; it has similarities to semiconductor-based demultiplexer schemes that connect binary addresses to devices or signals. As a proof of concept, we designed a scaffold structure consisting of six barcode subunits (‘bit positions’), each of which can have eight possible states as determined by the oligonucleotide sequence (octal, or base-8). Random selection of one of these eight sequences in the six subunit positions resulted in 262,144 independent barcodes that are accessible with only 48 oligonucleotides **(Supplementary Table S9)**. Each oligonucleotide sequence consists of a barcode (20bp) flanked by two adapter sequences (20bp each) that are required for combinatorial assembly into a larger barcode structure. For applicability to nanopore sequencing, we designed the barcode sequences to be uniquely identifiable using unique k-mer sequences **(Methods)**.

**Figure 4.**
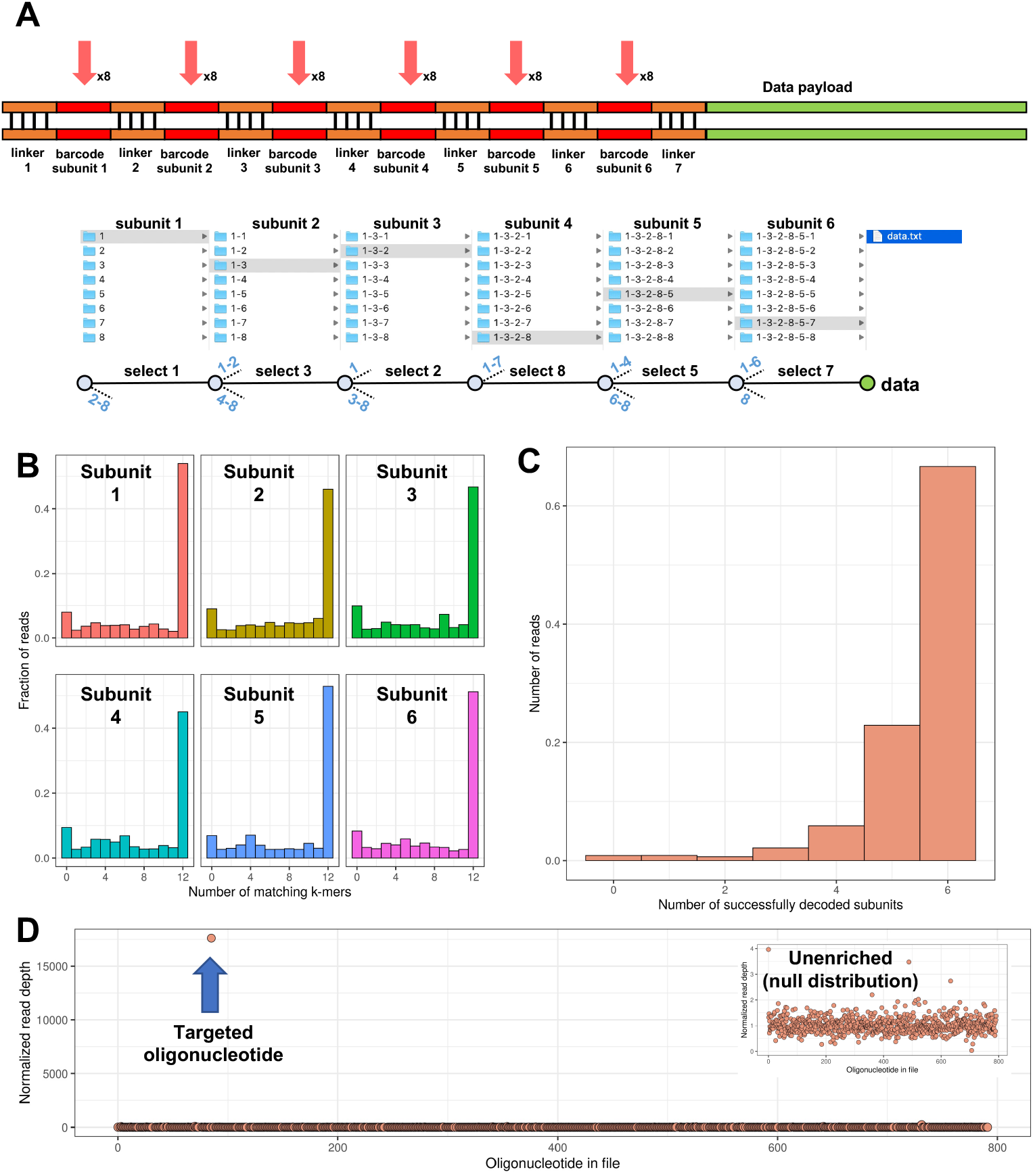
Massively parallel file barcoding with MDRAM. (A) Structure of combinatorial barcode addresses (CBA) for tagging data elements. A combinatorial barcode subunit and scaffold structure creates unique indexes for retrieving data elements. This creates a prototype hierarchical filesystem structure whereby individual data elements can be accessed by using primers sequentially targeting individual subunit locations. Bottom: a mock system relating CBA random access to filesystem tree traversal. (B) CBA sequence determination with nanopore sequencing. A k-mer matching scheme was used to determine the CBA sequence without alignment. The number of matching k-mers to the best match is shown for each subunit location. (C) The number of CBA sequences with perfect matches (no errors) is shown. (D) Targeting of a specific CBA to enrich for an oligonucleotide of interest. The median-normalized read count is plotted for each oligonucleotide for File 5 in Pool B. The arrow indicates the oligonucleotide of interest. Inset: The null distribution from an unenriched (non-targeted) pool of CBAs for File 5.

CBAs enable scalable random access of individual data elements. Random access is performed using a series of amplification steps with a small number of different primers. This sequential process enables traversal through the file hierarchical tree and reading specific DNA elements of interest **(Supplementary Figure S6)**. Initially, the DNA template consists of a pool of CBA-data element structures. To access an arbitrary data element, six sequential amplification reactions targeting each barcode subunit are performed, where the product of one reaction is diluted then used as the template for the next. Each amplification reaction traverses one subunit level of the CBA structure. The product from one amplification reaction is purified, diluted, and used as template for the next reaction. After six targeting reactions, the final amplicon belonging to the CBA-data element payload of interest is read through sequencing. For example, a primer corresponding to a barcode on subunit 1 was used in an amplification reaction. This amplifies sequences containing that barcode at subunit 1, resulting in enrichment of the nested elements in the hierarchical tree. The amplification product was purified and used for the template for the second amplification reaction with another primer corresponding to subunit 2, and so forth. Eventually, amplification for a sequence at barcode subunit 6 results in the data element of interest. All primers are listed in **Supplementary Table S9**.

To implement and validate CBA chemistry, we tagged individual oligonucleotides belonging to File 5 from Pool B with random CBAs. We amplified File 5 from Pool B and used the amplified DNA for random assignment of CBA addresses across the represented amplicons. We performed a one-step Gibson Assembly (*25*) reaction between amplicons of File 5 and a pool of all CBA scaffolding oligonucleotides **(Supplementary Table S9)** to generate a pool of CBAs, composed of randomly tagged oligonucleotides **(Methods)**. The CBA structure and the data elements are joined together by a linker sequence **(Fig. 4A, Supplementary Table S9)**. After the Gibson Assembly reaction, the products were amplified, cloned into topoisomerase-I activated vectors and transformed into *E. coli* to control for the overall library diversity.

To determine how individual oligonucleotides were associated with a given CBA, we nanopore sequenced the PCR products inserted into the vector. Lawns of bacterial colonies were pooled, amplified by PCR, and sequenced to determine the oligonucleotide data payload for each CBA. To perform address decoding, we used a k-mer matching algorithm that searches for unique sequencing that belong to each possible barcode at each subunit position. For each read and subunit position, we measured the number of matching k-mers (k=9) to every possible designed barcode sequence. Overall, we observed that ∼30-40% of sequences in each subunit had a perfect match to the designed barcode sequence **(Fig. 4B)**. As each barcode sequence consists of k-mers that are uniquely distinguishable from one another, sequences that had a less than perfect match can still be resolved. Overall, the entire CBA was successfully decoded for ∼70% of sequenced reads containing a CBA tag **(Fig. 4C)** which translates to an unrecoverable failure rate of approximately 6% at each barcode subunit, assuming that the errors between barcode subunits are independent.

Having determined the CBA distribution, we incorporated the CBA-File 5 constructs in the MDRAM format. Access of individual data elements was then performed on MDRAM by targeting for a specific sequence on each sequential barcode subunit **(Fig. 4A)**. The first amplification reaction was performed using primers corresponding to the common flanking sequences **(Supplementary Table S4 and Supplementary Table S9)**, after which subsequent CBA traversal reactions were performed in aqueous solution **(Supplementary Figure S6)**.

From our prior sequencing analysis of the original CBA pool, we counted 92,388 unique CBAs that are randomly linked to oligonucleotides from File 5 in Pool B. This number was determined using an inclusion criteria where k-mer scores for all other candidate barcodes for a given subunit must be zero. Using these CBA identities, we traversed the hierarchical tree with recombinase-based isothermal amplification (*26*) **(Methods)**. The targeting process involved six sequential amplification reactions – each amplification step uses a separate primer corresponding to the barcode sequence 5-11-23-29-39-48 and the reverse primer for File 5 **(Supplementary Table S9)**. After completion of these serial amplification steps, we sequenced the resulting amplicon. There were 17,600-fold more reads corresponding to the targeted oligonucleotide of interest compared the median coverage across all other File 5 oligonucleotides **(Fig. 4D)** and an approximately 10,900-fold enrichment versus the null unenriched distribution **(Fig. 4D, inset)**. Through this efficient and accurate targeting process, our results show a promising proof-of-concept for exponentially scalable random access of data elements on the MDRAM platform.

## DISCUSSION

In this work we developed MDRAM which is an end-to-end framework for storage of DNA data elements. This DNA data storage method features high-fidelity and repeated access of synthesized oligonucleotides, as well as efficient decoding of sequence features using a convolutional encoding/decoding scheme. This DNA data technology utilizes an efficient click chemistry scheme to attach synthesized DNA onto functionalized magnetic agarose beads. We demonstrated MDRAM’s capacity for conjugation of data-encoded oligonucleotides. Importantly, the chemistry itself is widely applicable for conjugation of other types of DNA onto the magnetic substrate – for example, we anticipate that longer strands of information-encoded DNA (*6*) or those from enzymatic synthesis (*27, 28*) would also be compatible with conjugation. Therefore, MDRAM is amenable for any type of biomolecule storage platform that is compatible with TCO-Tz labeling schemes.

We demonstrated that pools of data encoded oligonucleotides and their associated files can be selectively amplified from MDRAM over may iterations with little carryover contamination. Sequential read operations consisting of targeted PCR amplification of different DNA data files are performed on the same batch of magnetic beads with negligible amounts of sample loss. Aqueous solutions of DNA data files are consumed during any data readout operation and must be eventually exhausted or re-amplified. In contrast, MDRAM would potentially enable facile long-term repeated accessibility of data elements.

We demonstrated the data readout capability of MDRAM using an improved convolutional encoding/decoding workflow. When compared to other established methods, we found an overall reading cost improvement of approximately an order of magnitude. However, we note that a direct head-to-head comparison across methodologies is difficult due to various confounding factors, ranging from the use of different oligonucleotide synthesis providers, different amounts of data encoded, different oligonucleotide lengths synthesized and the ongoing improvements in nanopore sequencing technology. Some works utilized the now-discontinued 1D^2^ nanopore sequencing method to mitigate sequencing errors by reading the opposing strand (*7, 12*) but resulted in lower throughput compared to this work. Smaller file sizes were used in our study to focus on exploring multiple coding parameters and their impact on the overall reading cost. Smaller file sizes also effectively reduced the amount of indexing space within the synthesized oligonucleotides as compared to larger files. That being said, we anticipate has only a small effect on the overall reading and writing cost as the index size scales logarithmically with file size. Finally, we note that nanopore basecalling performance has improved dramatically, with current error rates of 4-5% compared to 10% as recently as 2019 (*29*), when other nanopore-based works were published (*7, 12*).

Our novel decoding workflow offers unique advantages over consensus-based schemes. By utilizing the rich soft information in the raw nanopore signal instead of working with basecalled reads, we sidestepped explicitly handling indel errors through consensus or an edit-distance code. The convolutional code is capable of correctly decoding a large fraction of reads without clustering or consensus, leading to an order-of-magnitude reduction in required coverage. Our resulting nanopore-based reading cost demonstrate a high level performance that matches coding schemes based on high-accuracy Illumina sequencing platforms (*16*). This equivalent performance is striking given that there is an order of magnitude difference in error rates between nanopore and Illumina sequencing. The framework has flexibility, seeing that it can be integrated into other neural network architectures as nanopore basecalling improves over time. Overall, the replication experiments utilizing MDRAM clearly demonstrate the robustness of the framework in effectively handling experimental variation due to coverage variance, dropouts, and the base quality of the sequencing run. Data could be encoded using a user’s choice of parameters optimizing on either decoding speed, writing cost, or robustness to errors.

Modern data storage systems utilize filesystems for random access of individual data elements amongst millions of files if not more. Individual primer synthesis and data access with conventional PCR does not scale to these numbers. In this work, we also demonstrated in MDRAM a proof-of-concept of an exponentially scalable data addressing technology called CBAs. By virtue of exponential scaling of combinatorial oligonucleotide assembly, we demonstrated individual readout of data elements with combinations of targeting oligonucleotides that can fit onto less than one 96-well plate. CBAs can also be incorporated onto data payloads with pre-determined barcode identities as opposed to being randomly assembled in our study; this would obviate the requirement for bulk sequencing to generate the CBA lookup table but may necessitate the use of robotic liquid handling to operate at scale. This work demonstrates that it is possible to mimic a basic feature of conventional filesystems – scalable random read access – on a DNA-based platform. In fact, in combination with our reduced reliance on sequence consensus, MDRAM could potentially enable real-time decoding with nanopore sequencing. With careful choice of convolutional coding parameters (m=6, r=3/4 and L=1), we achieved decoding times of 0.25 seconds per read while still achieving a reading cost of ∼4.5 bases/bit for a writing cost of ∼1 bases/bit – which is a 5x lower reading cost than another reported consensus-based approach (18). We also include the decoding speed for other parameters in **Supplementary Table S10**. We note that the outer Reed Solomon code operates in blocks and hence the block size should be kept small enough to reduce decoding latency but large enough to achieve reasonable erasure (sequence dropout) protection. Through a large-scale robotic MDRAM-to-sequencer integration and optimization of bioinformatic pipelines, we believe that the appropriate set of coding parameters can be used along with real-time basecalling to enable close to real-time repeated access and decoding of data as would be idealized in DNA-based data storage systems.

## Funding

This research was supported by the National Science Foundation and the Semiconductor Research Council (award number 1807371) under the SemiSynBio program. This work was also supported by a Beckman Technology Development Seed Grant.

## Author Contributions

B.T.L. and S.C. equally contributed to this work. B.T.L, S.C., T.W., and H.P.J. conceived and designed the study. B.T.L. and S.R. performed the experiments. S.C. and K.T. implemented the information encoding and decoding pipeline. B.T.L. and S.C. analyzed the data. M.W. provided feedback on the study design. T.W., and H.P.J. supervised the project. All authors contributed to the manuscript writing. We also thank Peter Griffin at the Stanford Genome Technology Center for helpful comments on the manuscript.

## Competing Interests

Authors declare that they have no competing interests.

## Data Availability

Source code for generating oligonucleotide sequences and analyzing sequencing reads derived from our convolutional coding scheme can be found here: https://github.com/shubhamchandak94/nanopore_dna_storage/tree/bonito. Sequence reads are deposited to NCBI’s Sequence Read Archive at PRJNA758230.

## LIST OF SUPPLEMENTARY MATERIALS

### Materials and Methods

Supplementary Text. Additional convolutional code optimization strategies.

Supplementary Figure S1. Convolutional inner code encoding.

Supplementary Figure S2. Convolutional inner code decoding.

Supplementary Figure S3. Outer coding strategy.

Supplementary Figure S4. Writing vs. reading cost across values of convolutional code memory (m) and rate (r).

Supplementary Figure S5. Writing vs. reading cost across values of convolutional code memory (m) and rate (r), for the one-CRC and two-CRC strategies.

Supplementary Figure S6. CBA traversal with sequential amplification reactions

Supplementary Table S1. Sequencing Metrics (Illumina)

Supplementary Table S2: Sequencing Metrics (Nanopore)

Supplementary Table S3: Encoding parameters for each synthetic DNA subpool in Pool B

Supplementary Table S4. Primer sequences for targeting Pool B

Supplementary Table S5. Percent of reads decoded correctly (list size 8 for decoding) across values of convolutional code memory (m) and rate (r).

Supplementary Table S6. Impact of improved basecalling pipeline and improved barcode removal on the percent of reads decoded correctly.

Supplementary Table S7. Nanopore decoding results.

Supplementary Table S8: Comparison to other platforms

Supplementary Table S9. CBA sequences

Supplementary Table S10. Decoding speed.

Supplementary Data S1. List of oligonucleotides for Pool A.

Supplementary Data S2. List of oligonucleotides for Pool B.

## Notes

### Competing Interest Statement

The authors have declared no competing interest.

