## Supplementary Materials for "Magnetic DNA random access memory with nanopore readouts and exponentially-scaled combinatorial addressing"

**This PDF file includes:**

Materials and Methods

Supplementary Text

Supplementary Figs. S1 to S6

Supplementary Tables S1 to S10

**Other Supplementary Materials for this manuscript include the following:**

Supplementary Data S1. List of oligonucleotides for Pool A.

Supplementary Data S2. List of oligonucleotides for Pool B.

### MATERIALS AND METHODS

#### Synthetic DNA samples (data files), and primers

We encoded a compressed data file of size 12.7 KB. Pool A and B are oligonucleotide pools that encode tarred, compressed and encrypted text containing a variety of poems, speeches such as the Gettysburg Address, lyrics and a declaration about human rights. The files include:

- gettysburg.txt: The Gettysburg Address by Abraham Lincoln
- mlkjr.txt: “I have a Dream” by Martin Luther King, Jr.
- rickastley.txt: Lyrics of “Never Gonna Give You Up” by Rick Astley
- poems.txt: A collection of poems:
    - “The Road Not Taken” by Robert Frost
    - “Stopping by Woods on a Snowy Evening” by Robert Frost
    - “If” by Rudyard Kipling
    - “Fire and Ice” by Robert Frost
    - “The Tyger” by William Blake
    - “A Psalm of Life” by Henry Wadsworth Longfellow
    - “Sonnet 18: Shall I compare thee to a summer’s day?” by William Shakespeare
    - “I Wandered Lonely as a Cloud” by William Wordsworth
    - “To Autumn” by John Keats
    - “Abou Ben Adhem” by Leigh Hunt
    - “Gitanjali 35” by Rabindranath Tagore
    - “Sympathy” by Paul Laurence Dunbar
    - “Caged Bird” by Maya Angelou
    - “Nothing Gold Can Stay” by Robert Frost
    - “The Rime of the Ancient Mariner” (excerpt) by Samuel Taylor Coleridge
- unhr.txt: The Universal Declaration of Human Rights

Synthetic DNA was prepared with the parameters ‘m’ (Convolutional code memory), ‘r’ (Convolutional code rate) with Reed Solomon code redundancy **(Supplementary Table 3)**. The specific procedure to encode data is described later in the Methods in **‘DNA encoding using convolutional codes’**. DNA sequences were synthesized by CustomArray (Pool A) and Agilent (Pool B) as a pool of oligonucleotides **(Supplementary Data S1 and S2)**. For read operations by PCR, primer sequences of length 25 were added on both sides of oligonucleotide sequences of interest to enable single file targeting. All sequences are also available at the URL https://github.com/shubhamchandak94/nanopore_dna_storage/tree/bonito/.

**DNA conjugation to magnetic beads**

Synthetic DNA was quantified by Qubit fluorescence (Thermo Fisher Scientific, Waltham, MA). 70ng of Pool A and 100 nanomoles of Pool B was used for bead conjugation. The DNA pool was denatured in a total volume of 45ul in 1X terminal transferase buffer (New England Biolabs, Ipswich, MA) with 1X (5ul) $CoCl_2$ additive (New England Biolabs, Ipswich, MA) for 5 minutes at 95C followed by ramping to at 4C at 100% ramping rate. The tailing reactions were performed in 50ul volumes with 100uM of TCO-PEG4-dUTP (Jena Bioscience, Jena, Germany) and 1ul of terminal transferase enzyme (New England Biolabs, Ipswich, MA) for 1 hour at 37C. After 1 hour of incubation, the reaction was purified using 1.8X Ampure XP beads (Beckman, Coulter, Brea, CA) and was resuspended in elution buffer (10mM Tris-HCl pH 8.0, 0.05% Tween 20).

Methyltetrazine-functionalized crosslinked agarose beads were custom ordered from Cube Biotech (Germany). 50ul of tetrazine beads were pipetted into a new PCR strip tube and washed twice with 100ul wash buffer (10mM Tris-HCl pH 8.0, 0.05% Tween 20) with low retention pipette tips. Afterwards, purified TCO-tailed DNA library was added onto the magnetic beads and incubated overnight at room temperature in a rotating mixer to prevent the beads from settling. After incubation, unconjugated library fragments were measured by Qubit. The conjugated DNA samples on magnetic beads were washed with wash buffer. The conjugation supernatant and wash steps can be saved for qPCR quantification. The samples were maintained in a storage buffer (10mM Tris-HCl pH 8.0, 0.1% Tween 20, 50% glycerol) at -20C.

**Read access of files with PCR**

Files were retrieved with PCR amplification using separate barcodes used as primers at 1uM concentration in 1X KAPA HiFi HotStart ReadyMix (Roche). Random access was directly performed by PCR by combining the master mix solution with the DNA-conjugated MDRAM. Primer sequences to target Pool A were previously published (*17*), and primers to target Pool B are listed in Supplementary Table S2. Amplification conditions were 45 seconds in 98C, 18 cycles of 98C for 15 seconds, 60C for 60 seconds, 72C for 60 seconds, and 72C for 60 seconds before a final hold at 4C. A magnetic bead separator was used to remove the amplified products from the beads. The supernatant containing amplified DNA libraries were purified using 1.8X volume of Ampure XP beads. The DNA-conjugated beads were washed with twice with by incubation of wash buffer 2 (98% formamide, 10mM Tris-HCl pH 8.0, 0.5% SDS) at 65C with mixing, twice with wash buffer (10mM Tris-HCl pH 8.0, 0.05% Tween 20), and stored with storage buffer for further use.

We also developed another cleanup step to eliminate carryover DNA between the reading of different data files. In later split-pool experiments involving repeated access of the same file, we further performed a high-stringency wash step consisting of digestion of pooled MDRAM beads with 1ul each of Exonuclease I and Exonuclease III (New England Biolabs, Ipswich, MA) in NEBuffer 2.1 for 30 minutes at room temperature, followed by heat inactivation for 20 minutes at

65C. Exonuclease I and exonuclease III are nucleases that digest double-stranded and single-stranded DNA from the 3' end. Because the MDRAM system conjugates DNA to the magnetic beads at the 3' end through the addition of TCO-functionalized nucleotides, the template DNA is protected from digestion. Consequently, carryover DNA fragments from the previous amplification is digested while the conjugated DNA remains protected. When accessing the same file repeatedly, we observed that this enzymatic digestion step eliminated the buildup of amplicons over successive read iterations. These beads were then washed twice with wash buffer (10mM Tris-HCl pH 8.0, 0.05% Tween 20), after which the beads were stored in storage buffer at -20C.

To sequence the amplified DNA, we performed end-repair, a-tailing, and ligation of sequencing adapters to the samples. For Illumina sequencing, DNA samples were subjected to end-repair and A-tailing process for 30 minutes at 20C and then 65C for 30 minutes with the Kapa HyperPrep workflow (Roche) in a thermal cycler, after which ligation was performed for 10 minutes at room temperature using Illumina UD barcoding adapters when sequencing using the Illumina platform. The ligation reaction was purified using 0.8X volume of Ampure XP beads under standard protocols. In the next step the library was amplified in 50ul KAPA HiFi HotStart Readymix (Roche) with an Illumina primer mix containing P5 and P7 sequences. Amplification conditions were 45 seconds at 98C, 12 cycles of 98C for 15 seconds, 60C for 30 seconds, 72C for 30 seconds, and 72C for 1 minute before a final hold at 4C. Amplified libraries were purified using 1.8X volume of Ampure XP beads. A ll amplified and purified libraries were quantified with Qubit analyzer and was sequenced on an Illumina iSeq with 50pM loading concentration.

For sequencing with the Oxford Nanopore platform, end-repaired and a-tailed DNA from the Kapa HyperPrep workflow was directly ligated with the AMX nanopore sequencing adapters using the LSK109 sequencing kit (Oxford Nanopore Technologies). We used the Native Barcoding Kit from Oxford Nanopore Technologies for multiplexed experiments. The ligation reaction was purified using 0.8X Ampure XP beads under standard protocols, with the exception of washing the beads with SFB buffer (Oxford Nanopore Technologies) instead of 80% ethanol, and eluted in EB buffer (Oxford Nanopore Technologies). All of the library was then immediately sequenced with a MinION R9.4.1 flowcell on the MinION nanopore sequencer for 48 hours.

**Sequence data processing and alignment**

Illumina sequence reads were demultiplexed using bclfastq v2.20 (Illumina). Reads were aligned using bwa-mem v2.17 (*30*), using the designed oligonucleotide pools as a reference. Each entry of the reference corresponds to a single oligonucleotide that was synthesized, making up a total of over 10,000 entries per reference. To assess targeting efficiency, read counts aligning to each sequence was tabulated using samtools v1.10 (*31*).

Nanopore sequence reads were basecalled and demultiplexed using Guppy (Oxford Nanopore Technologies). Sequence alignment was performed using minimap2 v2.17 (*32*). The designed oligonucleotide sequences were used as the reference, making up a total of over 10,000 entries. To assess targeting efficiency, read counts aligning to each sequence was tabulated using samtools v1.10 (*31*).

**DNA encoding using convolutional codes**

For nanopore sequencing, we use the coding scheme from Chandak et al. (*17*) with significant improvements through integrated basecalling and decoding stages **(Supplementary Figure 1)**. We extended the pipeline to work with recent nanopore basecallers such as guppy and bonito.

In addition, we modified the read trimming mechanism for removing the adapter and barcode sequences. These improvements in the inner code decoding pipeline lead to substantial improvements in the accuracy, in some cases doubling the number of the reads that are decoded correctly. Another modification was related to the fact that the previous scheme discarded reads where the CRC check failed, hence wasting non-trivial fractions of reads for which the decoding produced messages were mostly correct but had a few localized errors. The presence of such cases can be ascribed to the bursty nature of the decoding errors for convolutional codes. Therefore, we modified the strategy to use two CRCs for each oligo, protecting the first and second halves of the read, respectively **(Supplementary Figure 3)**. During decoding, we used a message from the list as long as at least one half has been decoded correctly, leading to further reduction in the number of reads needed for successful decoding. However, we discovered that the two CRC strategy only led to marginal gains in practice (see **Supplementary Text**), and hence focused on the simpler one CRC strategy for this analysis.

Input files were compressed and encrypted before encoding so that the input to the encoder appears random and does not produce excessive homopolymers (e.g., GGGG) which causes previously observed synthesis and sequencing errors (*17*). The encoded DNA sequences (including the sequencing primers) were of length around <200nt and were synthesized by Agilent under their 'HiFi' synthesis method.

**Integrating Viterbi decoding with bonito basecaller probabilities**

We integrated data decoding into v0.1 of the Bonito basecaller (*21*). Bonito is based on connectionist temporal classification (CTC) (*33*), which is a technique commonly used in speech recognition to bypass the need for labeling each timestep in the audio to a phoneme/letter.

Bonito utilizes a convolutional neural network that downsamples the raw signal and transforms it into a sequence of state probabilities at each time step, where the states are A, C, G, T and "blank" ("b"). Blank is a special state that allows the length of the basecall to be lower than the length of the probability vector. This state probability vector was decoded into a basecall in two ways:

1. Greedy: Here the most probable state is chosen at each time step and then a collapse operation is performed wherein repeated states are collapsed and blanks are removed. E.g., AAAbCCbCCCGGbbbTTT would be collapsed to ACCGT.
2. Beam search: This method exploits the fact that a single basecalled sequence corresponds to multiple state sequences due to the collapsing operation and the actual probability is the sum of probabilities all such state sequences. A set (beam) of candidate basecall sequences is maintained at each step, and each of them is extended at the next step. The size of the beam is kept constant, eliminating the lowest scoring sequences at each step.

For decoding the convolutional codes, we follow the beam search strategy, which is compatible with the list decoding used in Chandak et al. (*17*) **(Supplementary Figure 2)**. For the Viterbi decoding, we have states corresponding to (convolutional code state, position in base sequence). At each time step, each state stores a list of L candidate message prefixes (where L = list size or beam size). Along with the message prefixes, its current score is also stored (more precisely, we store the score for the top L paths ending with blank and the top L paths ending with non-blank CTC state). At the next step, we add one character (CTC state), add (using logsumexp operation) the scores for all the ways new bits can be added to extend the message prefix and take the top ones to maintain the list of size L. At the end of the Viterbi decoding process, we report the L messages corresponding to the state (convolutional code state = end state, position in message = total length). Other details including puncturing for high rate codes are described in Chandak et al. (*17*). We found that a list size of 8 provides close to optimal reading cost and higher list sizes do not provide much benefit. The increase in reading cost for list sizes below 8 is relatively small and the list size should be chosen based on the tradeoff between reading cost and computational complexity.

**Barcode removal during decoding**

The barcode/primer sequence needs to be removed before the convolutional code decoding which only expects the convolutional codeword as the input. This was done by first performing the basecalling, identifying the ends of the barcodes and then using only the probability vector corresponding to the convolutional codeword for the decoding. While the decoder is resilient to small imprecisions (few bases) in the barcode removal, larger errors can lead to suboptimal performance. We improved the performance by introducing a trimming penalty parameter to ensure that trimmed barcodes are correctly detected, while still giving preference to complete barcode sequences in the reads. In addition, we also made some other adjustments including better handling of cases involving chimeric reads where the start barcode might be found relatively later in the read than is usually expected.

**Computation of writing/reading cost**

The writing cost for each experiment was computed as $(file\ size\ in\ bytes\ \times\ 8)/(\#oligos\ total\ \times\ oligo\ length)$ while the reading cost was computed as $(file\ size\ in\ bytes\ \times\ 8)/(\#reads\ for\ decoding\ \times\ oligo\ length)$. In both cases, the oligo length did not include the primer sequence length. For the works that report coding density or bits per base excluding primers, the writing cost was computed as the reciprocal of these quantities. For works that report minimum coverage required for successful decoding, we computed the reading cost as $coverage \times\ writing\ cost$.

**Design and generation of exponential-scale combinatorial-barcode addresses (CBAs)**

We generated exponentially large combinations of unique barcode sequences using an established isothermal gene assembly method called Gibson Assembly (*25*). Each barcode subunit is flanked by common sequences that is required for the joining of oligonucleotides via this enzymatic assembly method. To generate these barcode sequences we randomly generated a set of 20-mer sequences that did not share any 9-mer sequences with each other. Forty eight of these sequences were chosen as the barcode sequences and seven chosen to be the flanking scaffold sequences. Oligonucleotides had the structure of, for example, [scaffold1]-[barcode1]-[scaffold2]. These oligonucleotides were synthesized by IDT without HPLC or PAGE purification and pooled together. The sequences are listed in **Supplementary Table S9**.

To link data elements to CBAs, we amplified files from Pool B where one of the primers contained a 5' CBA linking sequence (ATACGTTAGTTCGGCAGTAT) corresponding to the last scaffold sequence of the CBA and a forward primer of interest (eg. File 5). Amplification was performed in 1X NEBNext Ultra II Q5 Master Mix (New England Biolabs, Ipswich, MA), 1uM forward and reverse primer for the file, and 100 nanomoles of Pool B. Ramping conditions were 45 seconds at 98C, 18 cycles of 98C for 15 seconds, 60C for 60 seconds, 72C for 60 seconds, and 72C for 60 seconds before a final hold at 4C. Amplicons were purified with 1.8X Ampure XP beads with standard protocols. We then performed Gibson Assembly using 1X NEBuilder HiFi DNA Assembly Master Mix (New England Biolabs, Ipswich, MA), 1 picomole of amplicon product, and 10 picomoles of the CBA oligonucleotide pool. The reaction was incubated at 50C overnight, after which it was purified with 1.8X Ampure XP beads. To enrich for full-length products, we performed a second round of PCR in 1X NEBNext Ultra II Q5 Master Mix using 1uM flanking scaffolding primers (AGTGGAGTTCTCCGCATCAA and a reverse primer from a file in Pool B) with ramping conditions of 45 seconds in 98C, 18 cycles of 98C for 15 seconds, 60C for 60 seconds, 72C for 60 seconds, and 72C for 60 seconds before a final hold at 4C. These constructs were then purified with 1X Ampure XP beads under standard protocols. These amplicons were full length CBA-data structures whereby the data payload is an oligonucleotide sequence drawn from the array-synthesized oligonucleotide pool.

To control for the total library diversity, we performed TOPO cloning of the CBA-data constructs using the Zero Blunt TOPO PCR cloning kit (Thermo Fisher Scientific). TOPO-ligated vectors were then purified with 1.8X Ampure XP beads, eluted with water, electroporated into ElectroMAX DH10B *E. coli* cells (Thermo Fisher Scientific) and plated as a lawn onto Kanamycin-50 LB agar plates overnight at 37C. The next day, the lawn of cells were washed off with 2ml LB broth. 10ul of cells were diluted into 100ul elution buffer (10mM Tris-HCl pH 8.0, 0.05% Tween 20) and then lysed by incubating at 95C for 5 minutes. One ul of crude lysate was then amplified with 1X NEBNext Ultra II Q5 Master Mix using 1uM flanking scaffolding primers with ramping conditions of 45 seconds in 98C, 18 cycles of 98C for 15 seconds, 60C for 60 seconds, 72C for 60 seconds and 72C for 60 seconds before a final hold at 4C. MDRAM conjugation to magnetic beads is performed as above using 70ng of DNA.

**Random access of CBAs**

We first performed PCR on the MDRAM beads using 1uM primers consisting of a common flanking primer (AGTGGAGTTCTCCGCATCAA) corresponding to the first scaffold sequence and a corresponding reverse primer to File 5 in **Supplementary Table 4**. The beads were amplified with 1X NEBNext Ultra II Q5 Master Mix with ramping conditions of 45 seconds in 98C, 18 cycles of 98C for 15 seconds, 60C for 60 seconds, 72C for 60 seconds, and 72C for 60 seconds before a final hold at 4C. The supernatant was removed from the beads and each well was purified with 1.8X Ampure XP beads under standard conditions. The MDRAM beads were then washed with Exonuclease I and Exonuclease III as described above. The ground truth CBA and data payload information was generated by sequencing each amplicon on a R9 flowcell on the PromethION nanopore sequencer for 72 hours. Reads were basecalled into FASTQ files using Guppy, and reads were split into CBA and data payload sections using cutadapt (v2.9). CBA sequences were determined using a k-mer scanning algorithm that counts for unique k-mers found in each designed CBA sequence. They are matched with their corresponding data payload (oligonucleotide sequence) using the alignment program bwa. The inclusion criteria for CBAs in the ground truth lookup table was that for all six subunits there were no other k-mers associated with any other candidate barcodes.

We traversed the hierarchical tree of the CBA structure using a series of isothermal amplification steps with different primers. From the ground truth data, we chose a specific barcode sequence to target that corresponded to barcode sequence 5-11-23-29-39-48 tagged onto File 5. From the amplicons directly generated from the MDRAM beads, we performed multiple rounds of barcoded amplification using the barcode sequence **(Supplementary Table 4)** and the reverse primer targeting the entire file. Amplifications were in 1X TwistAmp Basic reaction mix (TwistDX, United Kingdom) with 1uM each primer, at 37C for 20 minutes followed by hold at 4C. The reaction mix was diluted five-fold before cleaning with 1.8X Ampure XP beads. The amplicon was then diluted 1:200 into the next amplification step traversing the hierarchical file tree. Sequencing libraries for the Oxford Nanopore platform were then generated using LSK110 chemistry. The amplicons were then sequenced using a R9 PromethION flowcell for 72 hours with data analysis as above.

**Supplementary Text. Additional convolutional code optimization strategies**

We explored additional optimization strategies to improve the performance of our convolutional coding scheme.

**Impact of additional CRCs**

We examined the impact of the new two-CRC strategy on the reading cost. Recall that the two-CRC strategy aimed at increasing the utilization of the convolutional decoded output by using parts of the read when the complete read is not decoded correctly. In **Supplementary Figure 5**, we observed that the results were mixed and there were no benefits of this strategy in improving data read quality. The reasons for this might include the fact that the overhead in reading cost for sequencing the second CRC offset any benefits in terms of fewer encoded reads per encoded sequence. Overall, this result pointed to the two-CRC strategy did not have an advantage over the simpler one-CRC strategy.

**Bonito model finetuning**

We performed finetuning (training) of the Bonito basecaller neural network parameters to exploit the fact that the DNA storage datasets differ significantly from the biological DNA datasets in terms of the strand length and base composition, among other factors. We generally followed the guidelines provided by the bonito tool usage guide, with some modifications. We first split the pool according to the convolutional code memory and rate (based on alignment) to obtain the training/validation split. To prepare the training dataset, the raw current reading sequences were first basecalled and aligned to the ground truth oligo sequences. The resulting pairs of raw current sequence and the oligo reference sequence formed the training data which is used to further train the pretrained model obtained from the Bonito basecaller (at a low learning rate to avoid overfitting). Some changes were made to the training pipeline:

- Reducing the chunk sizes during the training to be consistent with the oligo sequence lengths. Bonito typically uses chunk sizes of ~2000-4000 bases which is not suitable for this data since the oligo sequences are of length ~200.
- Modifying the evaluation metric to use global edit distance instead of local edit distance. Local edit distance metric computes edit distance between the read and the part of the oligo that the read aligns to, while the global edit distance is the edit distance between the read and the entire oligo. For our case, global edit distance is more suitable because we need accurate barcode detection for decoding, and training based on local edit distance leads to highly trimmed reads.
- Remove quality value and homopolymeric region filters while preparing the training data.

While we saw improved decoding accuracy on validation datasets within the same synthesis and sequencing pool, this did not generalize well and provided negligible improvements for the newly synthesized pool. Since the training did not generalize well to reads in different synthesis pools, we used the default bonito model for the final set of experiments.

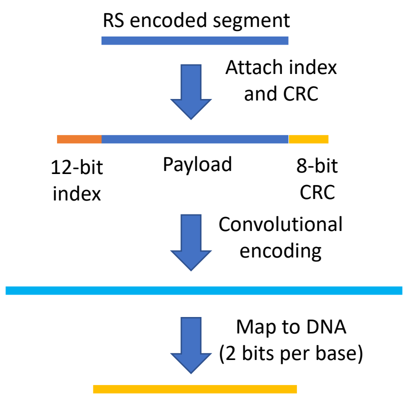


**Supplementary Figure S1. Convolutional inner code encoding.** Indexes and CRC are first attached to the RS segment from the encoder. Next, convolutional encoding is performed and is finally converted to a DNA sequence by performing the 2 bits per base mapping. Figure reproduced from (*17*).

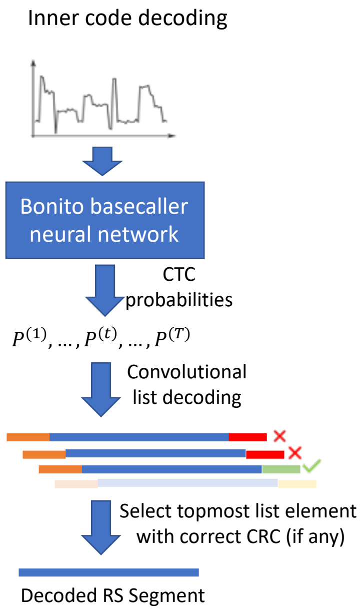


**Supplementary Figure S2. Convolutional inner code decoding.** The raw nanopore sequencing signal is passed through the bonito neural network to obtain the CTC probabilities, which are decoded using the Viterbi decoder which outputs a list of possible codewords. We select the highest scoring codeword that satisfies the CTC check, to obtain the decoded RS segment (to be used for outer code decoding later).

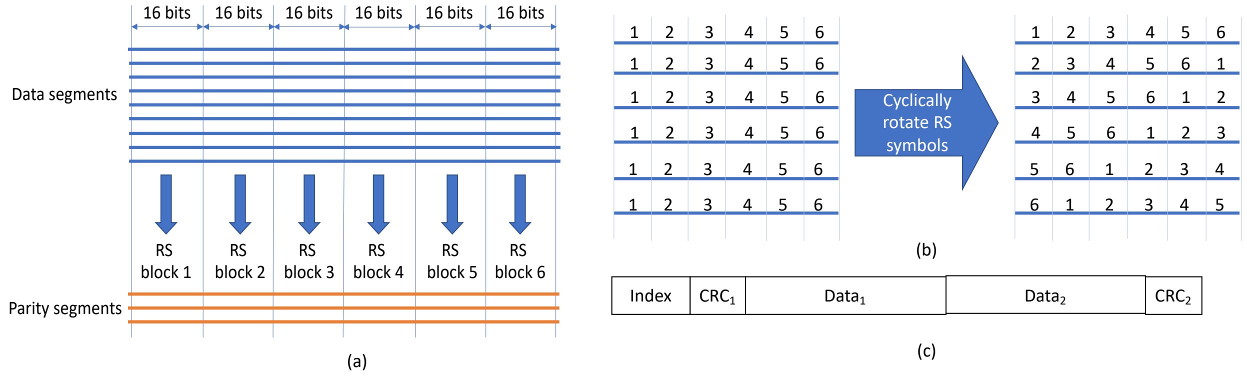


**Supplementary Figure S3. Outer coding strategy.** Two-CRC strategy for improving utilization of reads where only part of the read is correctly decoded. (a) RS encoding of the data based on Organick et al. (2018) and Chandak et al. (2020) (*7, 17*). In this example, each segment is split into six 16-bit RS symbols and the RS encoding is performed in a columnar manner to generate parity segments. The decoding succeeds as long as the number of erasures and errors in each column after inner code decoding is sufficiently small. (b) The RS symbols in each segment are cyclically rotated in a predefined order for reasons explained later. (c) The input to the convolutional code. The first CRC protects index+$data_1$, while the second CRC protects index+$data_2$. Based on the observation that in sequences with correct index, $data_1$ is more likely to be correct (due to the bursty error characteristics), we perform cyclic rotation of the RS symbols in the segments to make sure that each RS block has roughly balanced number of correct symbols for decoding. Note that the decoding succeeds once we obtain sufficiently many correctly decoded symbols from distinct oligos for each RS code (marked 1 through 6 in figure (b)). Using two CRC sequences enables partial utilization of the reads where part of the message is successfully decoded.

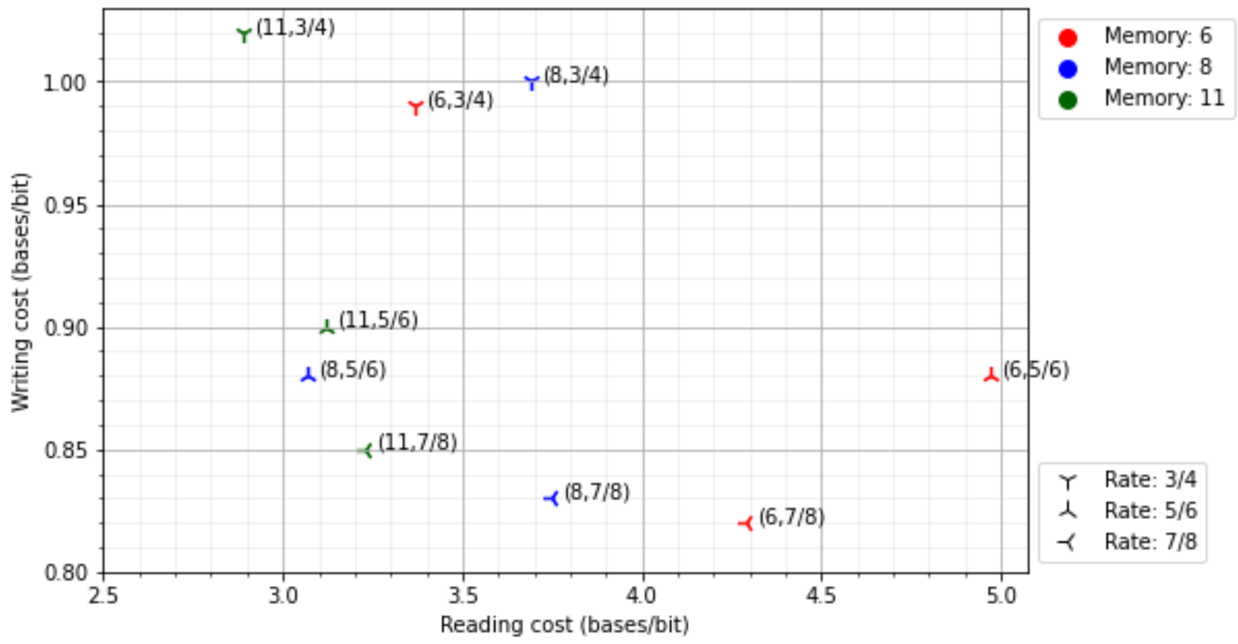


**Supplementary Figure S4: Writing vs. reading cost across values of convolutional code memory (m) and rate (r).** These results are for the subpools with the 1 CRC strategy. The small variation of writing cost with m (for fixed r) is due to the variation in the length of convolutional code padding bits.

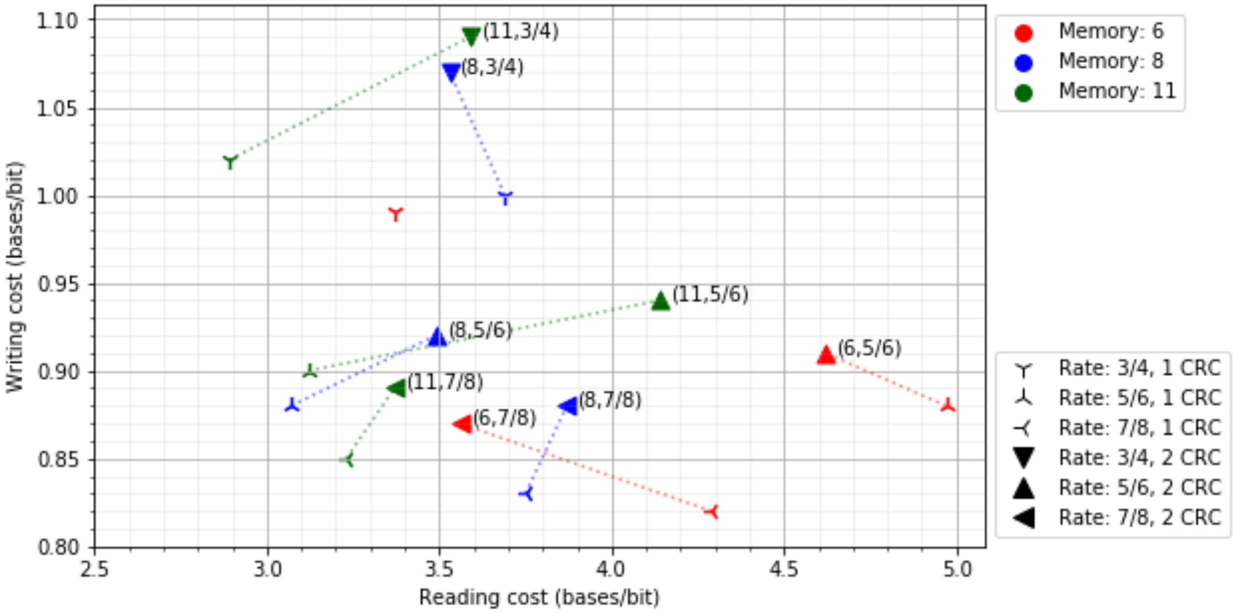


**Supplementary Figure S5. Writing vs. reading cost across values of convolutional code memory (m) and rate (r), for the one-CRC and two-CRC strategies.** The dotted lines connect the corresponding points for each (m,r) pair for clarity. The m=6, r=3/4, two-CRC subpool is not represented because of insufficient reads obtained in the amplification and sequencing process.

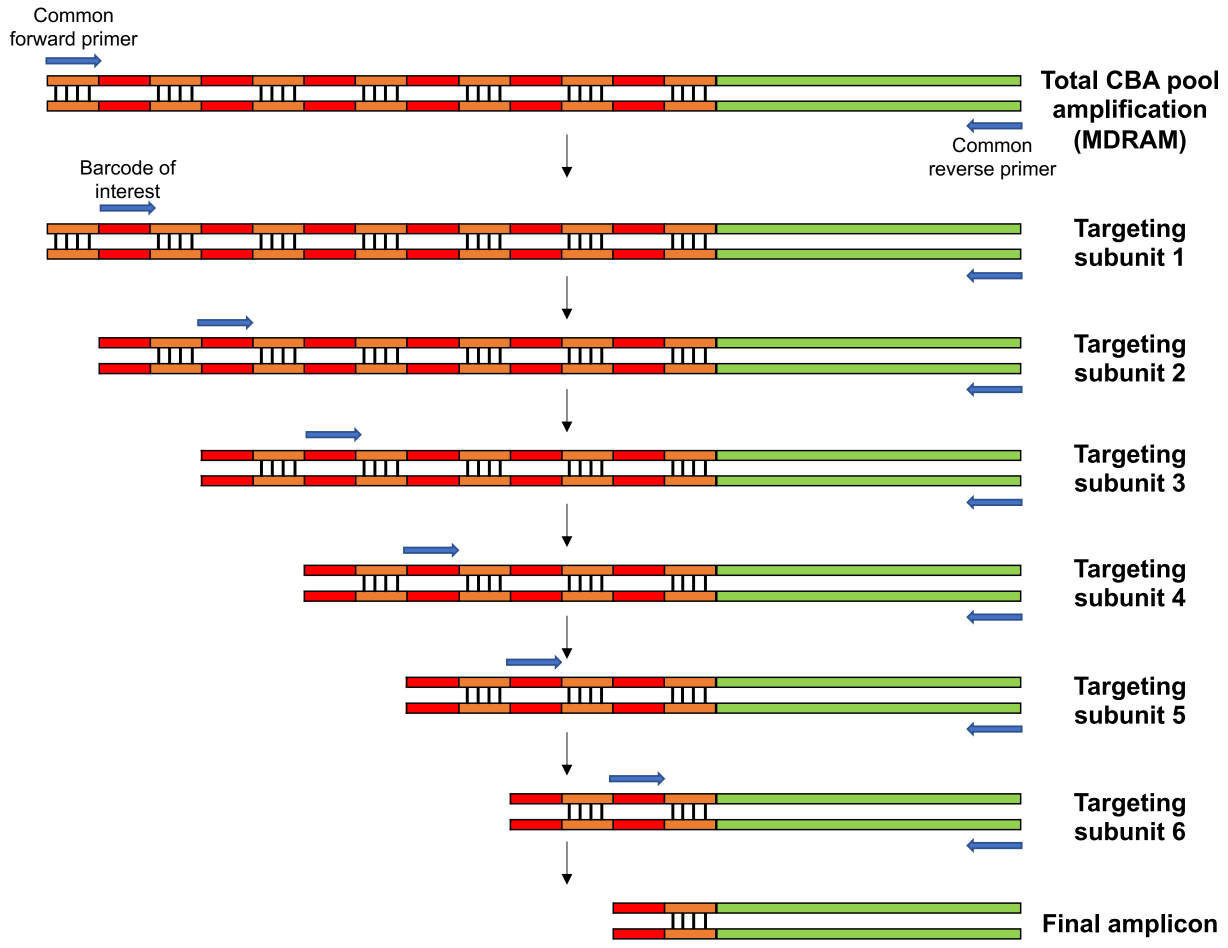


**Supplementary Figure S6. CBA traversal with sequential amplification reactions.** Primers corresponding to barcodes of interest are sequentially used in amplification reactions to traverse the CBA structure. Initially, the DNA template consists of a pool of CBA-data element structures. The product from one amplification reaction is purified, diluted, and used as template for the next reaction. Each amplification reaction traverses one subunit level of the CBA structure. After six targeting reactions, the final amplicon belonging to the CBA-data element payload of interest is retrieved. When used in the MDRAM format, total CBA pool amplification can be first performed by amplification using common flanking primers.

Supplementary Table S1. Sequencing Metrics (Illumina)

| Oligo Pool | File | Experiment | MDRAM-conjugated | Iteration | # Reads | % aligned |
|---|---|---|---|---|---|---|
| Pool A | 8 | Fig 2B,C | No | 1 | 323k | 90% |
| Pool A | 8 | Fig 2B - 1h | Yes | 1 | 562k | 89% |
| Pool A | 8 | Fig 2B,C - 24h | Yes | 1 | 348k | 66% |

Supplementary Table S2: Sequencing Metrics (Nanopore)

| Oligo Pool | File | MDRAM-Conjugated | Experiment | Iteration | # Reads | % aligned |
|---|---|---|---|---|---|---|
| Pool A | 1 | Yes | Fig 2D | 1 | 207.8k | 89% |
| Pool A | 2 | Yes | Fig 2D | 2 | 257.6k | 89% |
| Pool A | 3 | Yes | Fig 2D | 3 | 1044.9k | 89% |
| Pool A | 4 | Yes | Fig 2D | 4 | 451.0k | 88% |
| Pool A | 5 | Yes | Fig 2D | 5 | 257.3k | 90% |
| Pool A | 6 | Yes | Fig 2D | 6 | 283.9k | 91% |
| Pool A | 7 | Yes | Fig 2D | 7 | 78.9k | 80% |
| Pool A | 8 | Yes | Fig 2D | 8 | 665.0k | 81% |
| Pool A | 9 | Yes | Fig 2D | 9 | 625.6k | 89% |
| Pool A | 10 | Yes | Fig 2D | 10 | 481.0k | 89% |
| Pool A | 11 | Yes | Fig 2D | 11 | 152.0k | 88% |
| Pool A | 12 | Yes | Fig 2D | 12 | 504.2k | 89% |
| Pool A | 13 | Yes | Fig 2D | 13 | 832.9k | 87% |
| Pool B | 0,1,2,5,8 | No | Fig 3A,B | 1 | 9994.8k | 94% |
| Pool B | 0,1,2,5,8 | Yes | Fig 3C | 1 | 813.6k | 93% |
| Pool B | 0,1,2,5,8 | Yes | Fig 3C | 2 | 844.0k | 94% |
| Pool B | 0,1,2,5,8 | Yes | Fig 3C | 3 | 588.1k | 93% |
| Pool B | 0,1,2,5,8 | Yes | Fig 3C | 4 | 1813.0k | 93% |
| Pool B | 0,1,2,5,8 | Yes | Fig 3C | 5 | 1071.6k | 87% |
| Pool B | 0,1,2,5,8 | No | Fig 3C | 1 | 895.0k | 90% |
| Pool B | 5 | Yes - CBA | Fig 4B,C | 1 | 2449.2k | 91% |

Supplementary Table S3: Encoding parameters for each synthetic DNA subpool in Pool B

| Subpool # | m | r | Synthesis vendor | filesize (bytes) | # oligonucleotides (data) | # oligonucleotides (Reed Solomon) | # oligonucleotides (total) | bytes per oligo | CRC length | message length | oligonucleotide length |
|---|---|---|---|---|---|---|---|---|---|---|---|
| 0 | 6 | 0.75 | Agilent | 12672 | 704 | 176 | 880 | 18 | 8 | 165 | 114 |
| 1 | 8 | 0.75 | Agilent | 12672 | 704 | 176 | 880 | 18 | 8 | 164 | 115 |
| 2 | 11 | 0.75 | Agilent | 12672 | 704 | 176 | 880 | 18 | 8 | 164 | 117 |
| 3 | 6 | 0.83 | Agilent | 12672 | 634 | 158 | 792 | 20 | 8 | 180 | 112 |
| 4 | 8 | 0.83 | Agilent | 12672 | 634 | 158 | 792 | 20 | 8 | 180 | 113 |
| 5 | 11 | 0.83 | Agilent | 12672 | 634 | 158 | 792 | 20 | 8 | 180 | 115 |
| 6 | 6 | 0.88 | Agilent | 12672 | 576 | 144 | 720 | 22 | 8 | 197 | 116 |
| 7 | 8 | 0.88 | Agilent | 12672 | 576 | 144 | 720 | 22 | 8 | 196 | 117 |
| 8 | 11 | 0.88 | Agilent | 12672 | 576 | 144 | 720 | 22 | 8 | 197 | 119 |
| 9 | 6 | 0.75 | Agilent | 12672 | 792 | 198 | 990 | 16 | 8 | 156 | 108 |
| 10 | 8 | 0.75 | Agilent | 12672 | 792 | 198 | 990 | 16 | 8 | 157 | 110 |
| 11 | 11 | 0.75 | Agilent | 12672 | 792 | 198 | 990 | 16 | 8 | 157 | 112 |
| 12 | 6 | 0.83 | Agilent | 12672 | 634 | 158 | 792 | 20 | 8 | 189 | 117 |
| 13 | 8 | 0.83 | Agilent | 12672 | 634 | 158 | 792 | 20 | 8 | 188 | 118 |
| 14 | 11 | 0.83 | Agilent | 12672 | 634 | 158 | 792 | 20 | 8 | 189 | 120 |
| 15 | 6 | 0.88 | Agilent | 12672 | 634 | 158 | 792 | 20 | 8 | 188 | 111 |
| 16 | 8 | 0.88 | Agilent | 12672 | 634 | 158 | 792 | 20 | 8 | 188 | 112 |
| 17 | 11 | 0.88 | Agilent | 12672 | 634 | 158 | 792 | 20 | 8 | 188 | 114 |

Supplementary Table S4. Primer sequences for targeting Pool B

| File | Forward Primer (5'-3') | Reverse Primer (5'-3') |
|---|---|---|
| 0 | GCTACATGTATACTGCGAGACAGAC | GAGTGATGTGCGACTGCGACTATCG |
| 1 | TCTATCTACTCGTGCTCGCTAGCTG | ACATGCGACTAGTGCAGTGCAGACA |
| 2 | TGAGATCACAGCTACATAGTGAGAG | TGACTGCACACGACAGTGCTCTATC |
| 3 | AGCGTACACGACTGAGCACACTACG | TGCTAGCAGATGCGTGTGAGCGCAT |
| 4 | CATCAGCAGTAGAGAGTAGCGCGAT | GCAGACACTGCTAGCGTCGATGATA |
| 5 | GAGTCTCTAGCGCTACGAGATATAT | ACAGTCTGCTGATCTCATCAGAGCT |
| 6 | CACGAGATCTCAGTGTCGACACGTG | TAGCTGCGTCGAGATCACTATCACT |
| 7 | CGCTGCAGTCTATCTCTCTGTACAT | TCTCGCGAGATGTATCTCTACTAGC |
| 8 | GTGACTCTGCATATGCTGTCTCGAT | GTGCATCTCACGTGCACACACTCAG |
| 9 | ATGAGAGAGAGCGCTCTCTCATGAG | GTACTATGTACATCTATCGCGCTGT |
| 10 | GTCGTACTGTGACTCTCACTCGTGC | AGACATGTGTCGACACGTACGCAGA |
| 11 | CGCACGCATATCACGATGAGTAGCT | TGAGAGAGCTACTATATGTCTCGCA |
| 12 | CGTATGCTCGTATGAGCATAGAGTG | CATCACATCGATACATCTAGACGAC |
| 13 | GTGTGAGCTATGCGAGCGACGATCT | TGACGTACAGTGTACGACGTCGACG |
| 14 | TATGTACTAGCTAGTCACGCACACA | CGTGAGACTGACGACACAGCAGTGT |
| 15 | CTAGACGTGCGAGTATACTACTATG | TCACACGCGTCTGACTGCGTGTCAT |
| 16 | TGATCTATGATCGATGTACAGCGCG | AGTGTCAGAGATCGTGCGCGTGAGA |
| 17 | TCTCGCTCGAGCACAGAGATAGCGA | GCTCACAGCATCTACAGATACATGT |

Supplementary Table S5. Percent of reads decoded correctly (list size 8 for decoding) across values of convolutional code memory (m) and rate (r). These results are for the subpools comprising of a single CRC.

| **Convolutional code parameter** | **Memory m = 6** | **Memory m = 8** | **Memory m = 11** |
|---|---|---|---|
| **Rate r = 3/4** | 59.45% | 62.31% | 71.30% |
| **Rate r = 5/6** | 43.42% | 55.10% | 63.97% |
| **Rate r = 7/8** | 39.07% | 50.02% | 53.65% |

Supplementary Table S6. Impact of improved basecalling pipeline and improved barcode removal on the percent of reads decoded correctly (list size 8 for decoding). Flappie was used as the basecaller in Chandak et al. (2020), and the data from that study was used for these experiments. Guppy 3.4.5 is a basecaller from Oxford nanopore that is the production version of Flappie.

| Convolutional code memory (m) | Convolutional code rate (r) | Flappie | Guppy 3.4.5 | Bonito | Bonito + improved barcode removal |
|---|---|---|---|---|---|
| 8 | 1/2 | 68.93% | 72.52% | 74.20% | 87.10% |
| 8 | 3/4 | 26.90% | 31.87% | 34.40% | 58.90% |
| 8 | 5/6 | 21.67% | 26.91% | 37.10% | 39.60% |
| 11 | 1/2 | 62.80% | 65.90% | 62.90% | 88.30% |
| 11 | 3/4 | 39.62% | 45.13% | 58.30% | 67.70% |
| 11 | 5/6 | 25.91% | 33.02% | 44.30% | 53.40% |

Supplementary Table S7. Nanopore decoding results.

Supplementary Table 7A. Basecaller error rate (guppy 4.0.14)

| file# | m | r | Unconjugated | Iteration 1 | Iteration 2 | Iteration 3 | Iteration 4 | Iteration 5 |
|---|---|---|---|---|---|---|---|---|
| 0 | 6 | 0.75 | 5.36% | 5.37% | 5.31% | 5.54% | 5.35% | 5.46% |
| 1 | 8 | 0.75 | 5.58% | 5.63% | 5.60% | 5.85% | 5.61% | 5.82% |
| 2 | 11 | 0.75 | 5.79% | 5.87% | 5.79% | 5.98% | 5.95% | 5.95% |
| 5 | 11 | 0.83 | 5.65% | 5.64% | 5.59% | 5.85% | 5.69% | 5.72% |
| 8 | 11 | 0.88 | 5.72% | 5.86% | 5.76% | 5.94% | 5.78% | 6.02% |

Supplementary Table 7B. Coverage analysis (at 5x coverage subsampling): Normalized coverage variance - normalized by the variance for ideal Poisson sampling

| file# | m | r | Unconjugated | Iteration 1 | Iteration 2 | Iteration 3 | Iteration 4 | Iteration 5 |
|---|---|---|---|---|---|---|---|---|
| 0 | 6 | 0.75 | 1.32 | 1.39 | 1.57 | 1.60 | 1.72 | 1.77 |
| 1 | 8 | 0.75 | 1.33 | 1.45 | 1.59 | 2.74 | 15.90 | 19.38 |
| 2 | 11 | 0.75 | 1.47 | 1.64 | 1.50 | 1.49 | 1.62 | 1.89 |
| 5 | 11 | 0.83 | 1.90 | 1.86 | 1.86 | 1.76 | 1.79 | 2.00 |
| 8 | 11 | 0.88 | 1.28 | 1.26 | 1.55 | 1.49 | 1.83 | 2.01 |

Supplementary Table 7C. Fraction of oligos with 0 reads (at 5x coverage subsampling)

| file# | m | r | Unconjugated | Iteration 1 | Iteration 2 | Iteration 3 | Iteration 4 | Iteration 5 |
|---|---|---|---|---|---|---|---|---|
| 0 | 6 | 0.75 | 0.016 | 0.017 | 0.016 | 0.018 | 0.020 | 0.022 |
| 1 | 8 | 0.75 | 0.012 | 0.019 | 0.016 | 0.031 | 0.031 | 0.034 |
| 2 | 11 | 0.75 | 0.019 | 0.022 | 0.014 | 0.016 | 0.018 | 0.028 |
| 5 | 11 | 0.83 | 0.028 | 0.016 | 0.037 | 0.032 | 0.024 | 0.066 |
| 8 | 11 | 0.88 | 0.006 | 0.010 | 0.014 | 0.019 | 0.018 | 0.018 |

Supplementary Table 7D. Reading cost (success in 10 trials, number of reads used incremented in steps of 250)

| file# | m | r | Unconjugated | Iteration 1 | Iteration 2 | Iteration 3 | Iteration 4 | Iteration 5 |
|---|---|---|---|---|---|---|---|---|
| 0 | 6 | 0.75 | 3.65 | 3.65 | 3.65 | 3.94 | 3.94 | 4.50 |
| 1 | 8 | 0.75 | 3.97 | 3.97 | 4.54 | 4.25 | 4.54 | 5.39 |
| 2 | 11 | 0.75 | 3.17 | 3.46 | 3.17 | 3.17 | 3.46 | 3.75 |
| 5 | 11 | 0.83 | 3.69 | 3.69 | 3.97 | 3.69 | 3.40 | 4.25 |
| 8 | 11 | 0.88 | 4.40 | 4.40 | 4.70 | 4.70 | 4.70 | 5.58 |

Supplementary Table 7E. Number of reads for decoding (success in 10 trials, number of reads used incremented in steps of 250)

| file# | m | r | Unconjugated | Iteration 1 | Iteration 2 | Iteration 3 | Iteration 4 | Iteration 5 |
|---|---|---|---|---|---|---|---|---|
| 0 | 6 | 0.75 | 3250 | 3250 | 3250 | 3500 | 3500 | 4000 |
| 1 | 8 | 0.75 | 3500 | 3500 | 4000 | 3750 | 4000 | 4750 |
| 2 | 11 | 0.75 | 2750 | 3000 | 2750 | 2750 | 3000 | 3250 |
| 5 | 11 | 0.83 | 3250 | 3250 | 3500 | 3250 | 3000 | 3750 |
| 8 | 11 | 0.88 | 3750 | 3750 | 4000 | 4000 | 4000 | 4750 |

Supplementary Table 7F. Percent correctly decoded

| file# | m | r | Unconjugated | Iteration 1 | Iteration 2 | Iteration 3 | Iteration 4 | Iteration 5 |
|---|---|---|---|---|---|---|---|---|
| 0 | 6 | 0.75 | 51.08% | 52.06% | 52.39% | 51.05% | 51.32% | 47.28% |
| 1 | 8 | 0.75 | 50.58% | 50.83% | 46.93% | 47.03% | 47.39% | 40.09% |
| 2 | 11 | 0.75 | 61.62% | 61.16% | 63.29% | 62.64% | 60.46% | 56.07% |
| 5 | 11 | 0.83 | 51.82% | 52.41% | 49.34% | 49.46% | 51.94% | 47.80% |
| 8 | 11 | 0.88 | 37.49% | 39.00% | 37.82% | 36.75% | 37.66% | 31.82% |

Supplementary Table S8. Comparison to other platforms

| Writing cost (bases/bit) | Reading cost (bases/bit) | Min Coverage | Technology | Paper | Experimental parameters |
|---|---|---|---|---|---|
| 0.65 | 6.8 | 10.46 | Illumina | Erlich and Zielinski (2017) | |
| 0.91 | 4.55 | 5 | Illumina | Organick et al. (2018) | |
| 0.91 | 31.39 | 34.49 | Nanopore | Organick et al. (2018) | |
| 0.91 | 20.02 | 22 | Nanopore | Lopez et al. (2019) | |
| 0.91 | 2.73 | 3 | Illumina | Chandak et al. (2019) | LDPC w/ 50% redundancy |
| 0.67 | 3.82 | 5.7 | Illumina | Chandak et al. (2019) | LDPC w/ 10% redundancy |
| 0.93 | 7.01 | 7.54 | Nanopore | Chandak et al. (2020) | m=11,r=5/6 |
| 1.07 | 6.59 | 3.56 | Nanopore | Chandak et al. (2020) | m=14,r=3/4 |
| 1.85 | 4.42 | 4.13 | Nanopore | Chandak et al. (2020) | m=14,r=1/2 |
| 0.99 | 3.65 | 3.69 | Nanopore | This work | m=6, r=3/4 |
| 1.00 | 3.97 | 3.98 | Nanopore | This work | m=8, r=3/4 |
| 1.02 | 3.17 | 3.13 | Nanopore | This work | m=11, r=3/4 |
| 0.90 | 3.69 | 4.10 | Nanopore | This work | m=11, r=5/6 |
| 0.85 | 4.40 | 5.21 | Nanopore | This work | m=11, r=7/8 |

Supplementary Table S9. CBA sequences

| Subunit | Oligonucleotide # | Barcode sequence | Oligonucleotide sequence (5'-3') |
|---|---|---|---|
| 1 | 1 | GGTATATAGTGTCTTCTTGA | AGTGGAGTTCTCCGCATCAAGGTATATAGTGTCTTCTTGATACTGTAGTGCTGCCTTAAT |
| 1 | 2 | GTGCCAAGAACGCACTAGGA | AGTGGAGTTCTCCGCATCAAGTGCCAAGAACGCACTAGGATACTGTAGTGCTGCCTTAAT |
| 1 | 3 | CTGGTCTCTTCGGTCTGGAT | AGTGGAGTTCTCCGCATCAACTGGTCTCTTCGGTCTGGATTACTGTAGTGCTGCCTTAAT |
| 1 | 4 | TCTACTAATCAAGCGTTCGT | AGTGGAGTTCTCCGCATCAATCTACTAATCAAGCGTTCGTTACTGTAGTGCTGCCTTAAT |
| 1 | 5 | CTCTAATGTCCGAGCACACA | AGTGGAGTTCTCCGCATCAACTCTAATGTCCGAGCACACATACTGTAGTGCTGCCTTAAT |
| 1 | 6 | CTCTCCACTGTGACAAGTTG | AGTGGAGTTCTCCGCATCAACTCTCCACTGTGACAAGTTGTACTGTAGTGCTGCCTTAAT |
| 1 | 7 | ATACGACATAACTTCCGCTG | AGTGGAGTTCTCCGCATCAAATACGACATAACTTCCGCTGTACTGTAGTGCTGCCTTAAT |
| 1 | 8 | ACTATAACTTAACGTGAGCG | AGTGGAGTTCTCCGCATCAAACTATAACTTAACGTGAGCGTACTGTAGTGCTGCCTTAAT |
| 2 | 9 | GGACAACGCCTTCTTCTCAA | TACTGTAGTGCTGCCTTAATGGACAACGCCTTCTTCTCAAAGTATTCATATTAAGACGAA |
| 2 | 10 | CAGGTACTGCGTCTATATGG | TACTGTAGTGCTGCCTTAATCAGGTACTGCGTCTATATGGAGTATTCATATTAAGACGAA |
| 2 | 11 | TACATCTGGAATGATCGTGA | TACTGTAGTGCTGCCTTAATTACATCTGGAATGATCGTGAAGTATTCATATTAAGACGAA |
| 2 | 12 | CGAAGGCAATCGACTAATCG | TACTGTAGTGCTGCCTTAATCGAAGGCAATCGACTAATCGAGTATTCATATTAAGACGAA |
| 2 | 13 | ACGTTATCGGTGCCTCGTAC | TACTGTAGTGCTGCCTTAATACGTTATCGGTGCCTCGTACAGTATTCATATTAAGACGAA |
| 2 | 14 | GGCGTTGTATCTTCCAGCAA | TACTGTAGTGCTGCCTTAATGGCGTTGTATCTTCCAGCAAAGTATTCATATTAAGACGAA |
| 2 | 15 | CGCATACGTGCTTACTCGAG | TACTGTAGTGCTGCCTTAATCGCATACGTGCTTACTCGAGAGTATTCATATTAAGACGAA |
| 2 | 16 | CTGCGTTACGCCAACACCAC | TACTGTAGTGCTGCCTTAATCTGCGTTACGCCAACACCACAGTATTCATATTAAGACGAA |
| 3 | 17 | GTCCAGGTAGAGCCTATGAG | AGTATTCATATTAAGACGAAGTCCAGGTAGAGCCTATGAGCAGGCAACCTCGCAACCTCT |
| 3 | 18 | TTGCATACTTCTTGATGCTT | AGTATTCATATTAAGACGAATTGCATACTTCTTGATGCTTCAGGCAACCTCGCAACCTCT |
| 3 | 19 | AGAACTCACTCGCTGATCGG | AGTATTCATATTAAGACGAAAGAACTCACTCGCTGATCGGCAGGCAACCTCGCAACCTCT |
| 3 | 20 | GTGGAAGTTATCGATGCGAA | AGTATTCATATTAAGACGAAGTGGAAGTTATCGATGCGAACAGGCAACCTCGCAACCTCT |
| 3 | 21 | CTCTATGAACTGTAACCGCC | AGTATTCATATTAAGACGAACTCTATGAACTGTAACCGCCCAGGCAACCTCGCAACCTCT |
| 3 | 22 | TTAATTCCATTGAGAGGTTC | AGTATTCATATTAAGACGAATTAATTCCATTGAGAGGTTCCAGGCAACCTCGCAACCTCT |
| 3 | 23 | TAGACCATATACAAGTCTGG | AGTATTCATATTAAGACGAATAGACCATATACAAGTCTGGCAGGCAACCTCGCAACCTCT |
| 3 | 24 | TCCGAAGCTTGGTGACTCTC | AGTATTCATATTAAGACGAATCCGAAGCTTGGTGACTCTCCAGGCAACCTCGCAACCTCT |
| 4 | 25 | GAATACCGAGCGATACAGAA | CAGGCAACCTCGCAACCTCTGAATACCGAGCGATACAGAAATCACACTAGTGAATCGTGA |
| 4 | 26 | AGGTGGACAGTAATAGTATG | CAGGCAACCTCGCAACCTCTAGGTGGACAGTAATAGTATGATCACACTAGTGAATCGTGA |
| 4 | 27 | GCAAGTACGACAGCCTATTC | CAGGCAACCTCGCAACCTCTGCAAGTACGACAGCCTATTCATCACACTAGTGAATCGTGA |
| 4 | 28 | AGAAGCTCACCTCAAGTGAC | CAGGCAACCTCGCAACCTCTAGAAGCTCACCTCAAGTGACATCACACTAGTGAATCGTGA |
| 4 | 29 | TAACTCCAAGGTGCGGTGGC | CAGGCAACCTCGCAACCTCTTAACTCCAAGGTGCGGTGGCATCACACTAGTGAATCGTGA |
| 4 | 30 | ACCAACGGAATGCTTACTGG | CAGGCAACCTCGCAACCTCTACCAACGGAATGCTTACTGGATCACACTAGTGAATCGTGA |
| 4 | 31 | TCCAAGTATCAGTCCTGGTC | CAGGCAACCTCGCAACCTCTTCCAAGTATCAGTCCTGGTCATCACACTAGTGAATCGTGA |
| 4 | 32 | GCTCGATCTGCTAGATCTAG | CAGGCAACCTCGCAACCTCTGCTCGATCTGCTAGATCTAGATCACACTAGTGAATCGTGA |
| 5 | 33 | AGGTTCACCTCTGGATACGC | ATCACACTAGTGAATCGTGAAGGTTCACCTCTGGATACGCAGGATTGCTGATGGATCTGG |
| 5 | 34 | AGACTTCGTGAGCATGCTGG | ATCACACTAGTGAATCGTGAAGACTTCGTGAGCATGCTGGAGGATTGCTGATGGATCTGG |
| 5 | 35 | TCTATCAGCACCTCTACACA | ATCACACTAGTGAATCGTGATCTATCAGCACCTCTACACAAGGATTGCTGATGGATCTGG |
| 5 | 36 | TGAATCTAGGTGGTTAACAG | ATCACACTAGTGAATCGTGATGAATCTAGGTGGTTAACAGAGGATTGCTGATGGATCTGG |
| 5 | 37 | TTGACGTATCTCATGATACC | ATCACACTAGTGAATCGTGATTGACGTATCTCATGATACCAGGATTGCTGATGGATCTGG |
| 5 | 38 | CGCTTCTTCCAGTCAGCATG | ATCACACTAGTGAATCGTGACGCTTCTTCCAGTCAGCATGAGGATTGCTGATGGATCTGG |
| 5 | 39 | CAGACGACCTATGCGAGAAT | ATCACACTAGTGAATCGTGACAGACGACCTATGCGAGAATAGGATTGCTGATGGATCTGG |
| 5 | 40 | TGAGCATTATGCCTCCTCGC | ATCACACTAGTGAATCGTGATGAGCATTATGCCTCCTCGCAGGATTGCTGATGGATCTGG |
| 6 | 41 | ACGACCTAGACTCACAGCCA | AGGATTGCTGATGGATCTGGACGACCTAGACTCACAGCCAATACGTTAGTTCGGCAGTAT |
| 6 | 42 | TCAACGTTGCTTGGAAGTTC | AGGATTGCTGATGGATCTGGTCAACGTTGCTTGGAAGTTCATACGTTAGTTCGGCAGTAT |
| 6 | 43 | AGTTGCTACCATCCTTCATG | AGGATTGCTGATGGATCTGGAGTTGCTACCATCCTTCATGATACGTTAGTTCGGCAGTAT |
| 6 | 44 | TGAACAACCTGGTGGACTGG | AGGATTGCTGATGGATCTGGTGAACAACCTGGTGGACTGGATACGTTAGTTCGGCAGTAT |
| 6 | 45 | GCGTAAGCGGTGATCGACGC | AGGATTGCTGATGGATCTGGGCGTAAGCGGTGATCGACGCATACGTTAGTTCGGCAGTAT |
| 6 | 46 | CTCATCTGATGGCAGTACCG | AGGATTGCTGATGGATCTGGCTCATCTGATGGCAGTACCGATACGTTAGTTCGGCAGTAT |
| 6 | 47 | TACACTGACTGTCAACCACT | AGGATTGCTGATGGATCTGGTACACTGACTGTCAACCACTATACGTTAGTTCGGCAGTAT |
| 6 | 48 | TCTTAAGATCTTACATAGTC | AGGATTGCTGATGGATCTGGTCTTAAGATCTTACATAGTCATACGTTAGTTCGGCAGTAT |

Supplementary Table S10. Decoding speed (on a Intel(R) Xeon(R) CPU E5-2630 v4 @ 2.20GHz processor).

| | | | Time per read (in s) on 1 thread | | Reading cost | |
|---|---|---|---|---|---|---|
| m | r | Writing cost | L=1 | L=8 | L=1 | L=8 |
| 6 | 3/4 | 0.990 | 0.25 | 2 | 4.217 | 3.374 |
| 8 | 3/4 | 0.998 | 1 | 8 | 4.254 | 3.687 |
| 11 | 3/4 | 1.016 | 8 | 64 | 3.174 | 2.885 |
| 6 | 5/6 | 0.875 | 0.25 | 2 | 6.353 | 4.972 |
| 8 | 5/6 | 0.883 | 1 | 8 | 3.623 | 3.065 |
| 11 | 5/6 | 0.898 | 8 | 64 | 3.687 | 3.120 |
| 6 | 7/8 | 0.824 | 0.25 | 2 | 5.435 | 4.291 |
| 8 | 7/8 | 0.831 | 1 | 8 | 4.328 | 3.751 |
| 11 | 7/8 | 0.845 | 8 | 64 | 3.522 | 3.228 |
